# Rapid, biochemical tagging of cellular activity history in vivo

**DOI:** 10.1101/2023.09.06.556431

**Authors:** Run Zhang, Maribel Anguiano, Isak K. Aarrestad, Sophia Lin, Joshua Chandra, Sruti S. Vadde, David E. Olson, Christina K. Kim

**Affiliations:** Biomedical Engineering Graduate Group, University of California, Davis, Davis, CA 95616; Center for Neuroscience, University of California, Davis, Davis, CA 95618; Neuroscience Graduate Group, University of California, Davis, Davis, CA 95618; Institute for Psychedelics and Neurotherapeutics, University of California, Davis, Davis, CA 95616; Department of Neurology, University of California, Davis, Sacramento, CA 95817; Department of Chemistry, University of California, Davis, Davis, CA 95616; Department of Biochemistry and Molecular Medicine, University of California, Davis, Sacramento, CA 95817

## Abstract

Intracellular calcium (Ca^2+^) is ubiquitous to cell signaling across all biology. While existing fluorescent sensors and reporters can detect activated cells with elevated Ca^2+^ levels, these approaches require implants to deliver light to deep tissue, precluding their noninvasive use in freely-behaving animals. Here we engineered an enzyme-catalyzed approach that rapidly and biochemically tags cells with elevated Ca^2+^ in vivo. Ca^2+^-activated Split-TurboID (CaST) labels activated cells within 10 minutes with an exogenously-delivered biotin molecule. The enzymatic signal increases with Ca^2+^ concentration and biotin labeling time, demonstrating that CaST is a time-gated integrator of total Ca^2+^ activity. Furthermore, the CaST read-out can be performed immediately after activity labeling, in contrast to transcriptional reporters that require hours to produce signal. These capabilities allowed us to apply CaST to tag prefrontal cortex neurons activated by psilocybin, and to correlate the CaST signal with psilocybin-induced head-twitch responses in untethered mice.

## MAIN TEXT

Dynamic changes in intracellular ion concentrations allow cells to respond and adapt to their local environment, ultimately contributing to the normal physiological functioning of organisms. For example, neurons, the basic functional units of the brain, can be activated by various external stimuli or pharmacological compounds, leading to rapid fluctuations in intracellular calcium ion (Ca^2+^) concentrations. Thus, activity among complex neural networks can be measured using cellular changes in Ca^2+^ levels as a direct proxy for neuronal firing. Genetically-encodable Ca^2+^ indicators have transformed our ability to record neural activity in awake and behaving animals^1–4^. However, a major limitation of fluorescent sensors is that their read-out is transient, and they generally require invasive methods to gain optical access to deep brain structures. This can make it challenging to couple the activity history of a given neuron with its numerous other cellular properties (e.g., precise spatial localization, RNA expression, or protein expression).

To overcome this issue, previous efforts have focused on designing orthogonal transcriptional reporters (FLARE^5^, FLiCRE^6^, Cal-Light^7^) or fluorescent proteins (CaMPARI^8^) that can stably mark activated cells undergoing high intracellular Ca^2+^. However, these approaches implement light-sensitive proteins that require blue^9^ or UV^10^ light to restrict the time window of activity labeling in cells. This requirement hampers their scalability in deep brain regions, or in body areas where fibers for light delivery cannot be implanted. Alternative stable tagging approaches include immediate early gene (IEG)-based transcriptional reporters (TRAP2^11^, and tetTag^12, 13^), which utilize a drug injection instead of light to gate the activity labeling window. However, while IEG activity has been linked to neural activity in many cell-types^14^, it is not nearly as universal a read-out as Ca^2+^ is. Furthermore, the slow onset of IEG expression limits the ability to immediately tag and identify neurons activated during a specific time window. This is compounded by the fact that all transcription-based activity reporters take several hours (∼6-18 hrs^6, 15^) before sufficient levels of the reporter protein can be detected. Thus, there is a need for a strategy that enables non-invasive and rapid activity-dependent labeling of cells.

We designed an activity-dependent enzyme that can attach a small, biochemical handle to activated cells exhibited high intracellular Ca^2+^. Our strategy was to re-engineer and re-purpose a proximity-labeling enzyme, split-TurboID^16^, to report increased intracellular Ca^2+^ in living cells by tagging proteins with an exogenously delivered biotin molecule. Proximity labeling enzymes such as split-TurboID^16^ (and its predecessors, BioID^17^ and TurboID^18^) have been traditionally used to biotinylate nearby, statically-present proteins for downstream enrichment and analysis over long periods (typically 4-7 days in vivo). But they have not been engineered to detect dynamic changes in intracellular ion concentrations. Our design, CaST (Ca^2+^-activated Split-TurboID), enzymatically tags activated neurons within brief, user-defined time windows of exogenous biotin delivery. The biotinylated proteins can then immediately be read-out using any existing method for biotin-detection. Since the biotin molecule is both cell- and blood-brain barrier-permeable^19, 20^, this facilitates its application in living organisms.

## RESULTS

### Design and optimization of a Ca^2+^-activated Split-TurboID

The basic CaST design tethers the Ca^2+^-binding protein calmodulin (CaM) and a CaM-binding synthetic peptide M13 variant to either inactive half of split-TurboID (**Fig. 1A**). We postulated that under high cytosolic Ca^2+^ concentrations, the CaM fragment would be recruited to M13, resulting in the reconstitution and activation of split-TurboID. Upon simultaneous biotin supplementation, the reconstituted split-TurboID would then biotinylate itself and nearby proteins^16^, in a Ca^2+^-dependent manner. High Ca^2+^ alone should not result in signal, as the endogenous biotin levels are too low to result in substantial protein biotinylation; and exogenous biotin alone should not result in signal, as the split-TurboID fragments remain separated and inactive (**Fig. 1B**). Thus, CaST can act as a coincidence detector of both exogenous biotin and high intracellular Ca^2+^.

**Figure 1.**
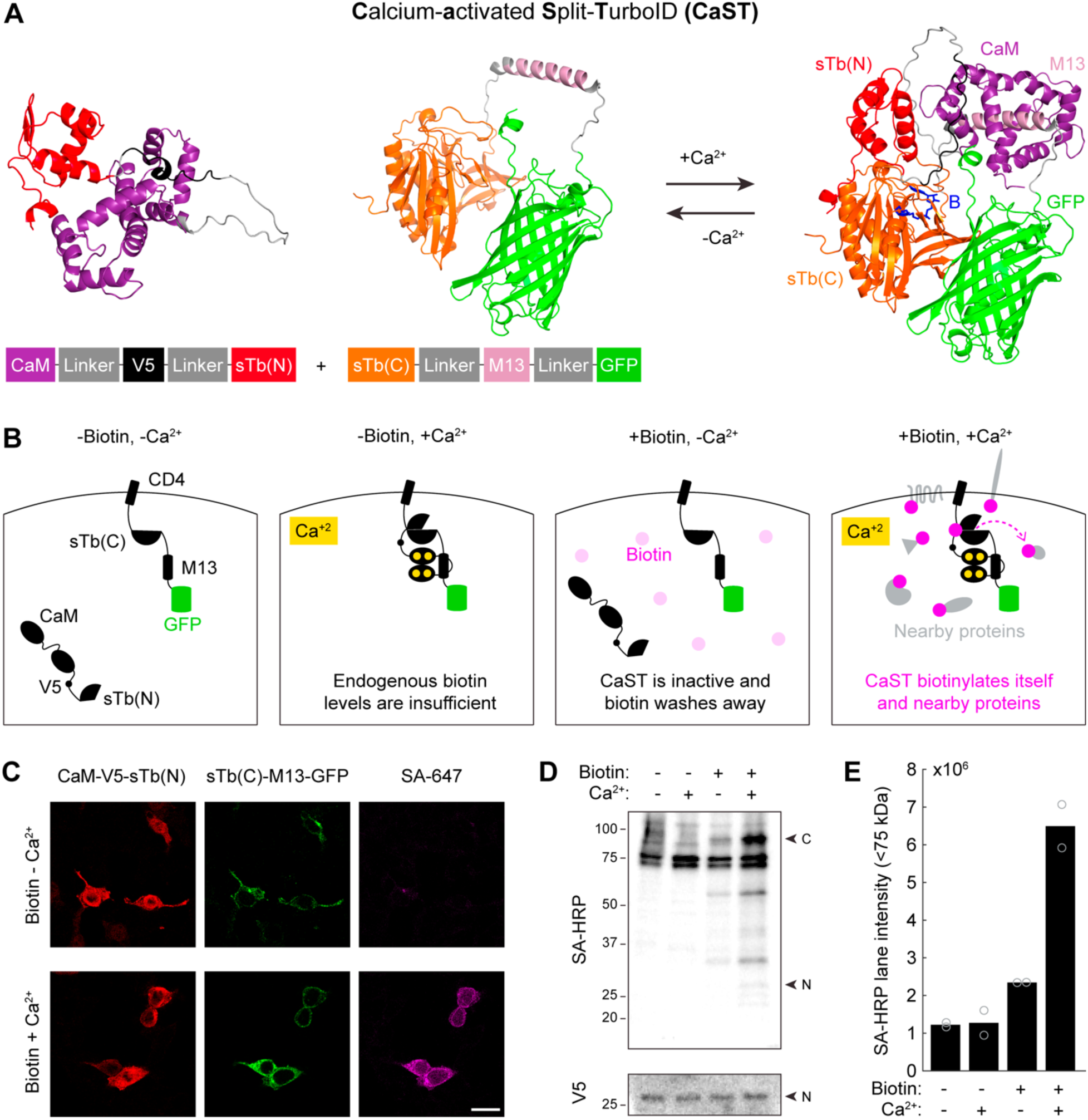
Design of a Ca^2+^-activated Split-TurboID (CaST). **A)** AlphaFold2^56, 57^ prediction of the protein structures for the two halves of CaST either in isolation (left; as expected in the absence of Ca^2+^), or in complex (right; as expected in the presence of high Ca^2+^). The two CaST components reversibly reconstitute in a Ca^2+^-dependent manner. The predicted biotin binding site is shown in blue. **B)** Schematic of CaST design for expression in HEK cells. The component with sTb(C)-M13- GFP is tethered to the membrane via the transmembrane domain of a cell membrane protein CD4^58^, while the CaM-V5 epitope tag^59^-sTb(N) component is expressed throughout the cytosol. CaST only tags proteins when cells are treated with biotin and exhibit elevated intracellular Ca^2+^. **C)** Example confocal images of HEK cells transfected with both components of CaST and treated with biotin ± Ca^2+^ for 30 minutes, as described in panel **C**. Cells were washed, fixed, and stained with anti-V5 and streptavidin-AlexaFluor647 (SA-647). The anti-V5 signal stains the CaM-sTb(N) component (left), while the GFP fluorescence shows the CD4-sTb(C)-M13 component (middle). Cells treated with high Ca^2+^ exhibit robust biotinylation of proteins, detected by SA-647 staining (right). Scale bar, 20 µm. **D)** HEK cells were transfected with CaST and treated with ± 50 µM biotin and ± Ca^2+^ (5 mM CaCl2 and 1 µM ionomycin) for 30 minutes. Cells were then washed with DPBS, and whole-cell lysates were collected and analyzed using a Western blot stained with streptavidin-Horse Radish Peroxidase (SA-HRP) or anti-V5/HRP. “N” indicates the expected size of the CaM-V5- sTb(N) fragment, while “C” indicates the expected size of the CD4-sTb(C)-M13-GFP fragment. **E)** Quantification of biotinylated proteins present in Western blot lanes of the experiment shown in panel **C**. Two independent biological replicates were quantified. The entire lane below the 75kDa endogenously biotinylated bands were included in the quantification (sum of the total raw intensity pixel values). A line plot profile spanning the entire blot is shown in **Extended Data** Figure 2.

Initially, we tested various approaches to tether CaM and M13 to either of the split-TurboID fragments, sTb(N) and sTb(C). Four different versions of the tool were transfected into HEK293T cells, with different conformational arrangements and subcellular localizations of the proteins (**Supplementary** Fig. 1A). Cells were treated with a combination of biotin with or without Ca^2+^ and an ionophore for 30 minutes, and then fixed and stained for biotinylated proteins using Streptavidin conjugated to Alexa-647 (SA-647). Both the GFP and SA-647 fluorescence were quantified for each cell and were ratioed to normalize for differences in expression levels across cells or experimental conditions (SA-647/GFP; see Methods). Importantly, all four versions had no SA-647 signal in the absence of exogenously delivered biotin. Of the four versions, we found that a membrane-tethered CD4-sTb(C)-M13-GFP with a cytosolic CaM-V5-sTb(N) resulted in the highest signal-to-background ratio (SBR) of biotin ± Ca^2+^ tagging (**Supplementary** Fig. 1B). We subsequently designated this optimized construct as our final CaST design (**Fig. 1B**). We also showed that a 5:2 transfection ratio of the two CaST fragments (CD4-sTb(C)-M13-GFP to CaM-V5-sTb(N)) yielded the highest SBR of all ratios tested (**Extended Data** Fig. 1A-C). Subsequent characterizations were performed using this optimal transfection ratio.

Immunohistochemistry and confocal imaging showed that both fragments of the tool are expressed in HEK cells, confirming that the low SA-647 signal in negative control conditions is not due to a lack of fragment co-expression (**Fig. 1C**). As expected, purposefully omitting either fragment of CaST in the presence of biotin and Ca^2+^ resulted in no biotinylation signal (**Extended Data** Fig. 1D). Western blot analysis confirmed that CaST-transfected HEK239T (CaST-HEK) cells treated with biotin and Ca^2+^ drove biotinylation across an array of nearby proteins, compared to negative control conditions (**Fig. 1D,E**, and **Extended Data** Fig. 2A-D).

### Characterization of CaST labeling in HEK cells

To further quantify the extent of biotin- and Ca^2+^-dependent CaST labeling, we performed fluorescence imaging across multiple fields of views (FOVs) in CaST-HEK cells treated with biotin ± Ca^2+^ (**Fig. 2A**). Single cell analysis confirmed that the Ca^2+^-dependent increase in SA- 647 labeling was present across cells with varying GFP expression levels, showing that it is not due to differences in expression levels of the tool across conditions (**Fig. 2B**). The distributions of normalized SA-647/GFP fluorescence also differed between cells treated with or without Ca^2+^ (**Fig. 2C**). These results demonstrate that CaST robustly detects elevated intracellular Ca^2+^ levels in living cells.

**Figure 2.**
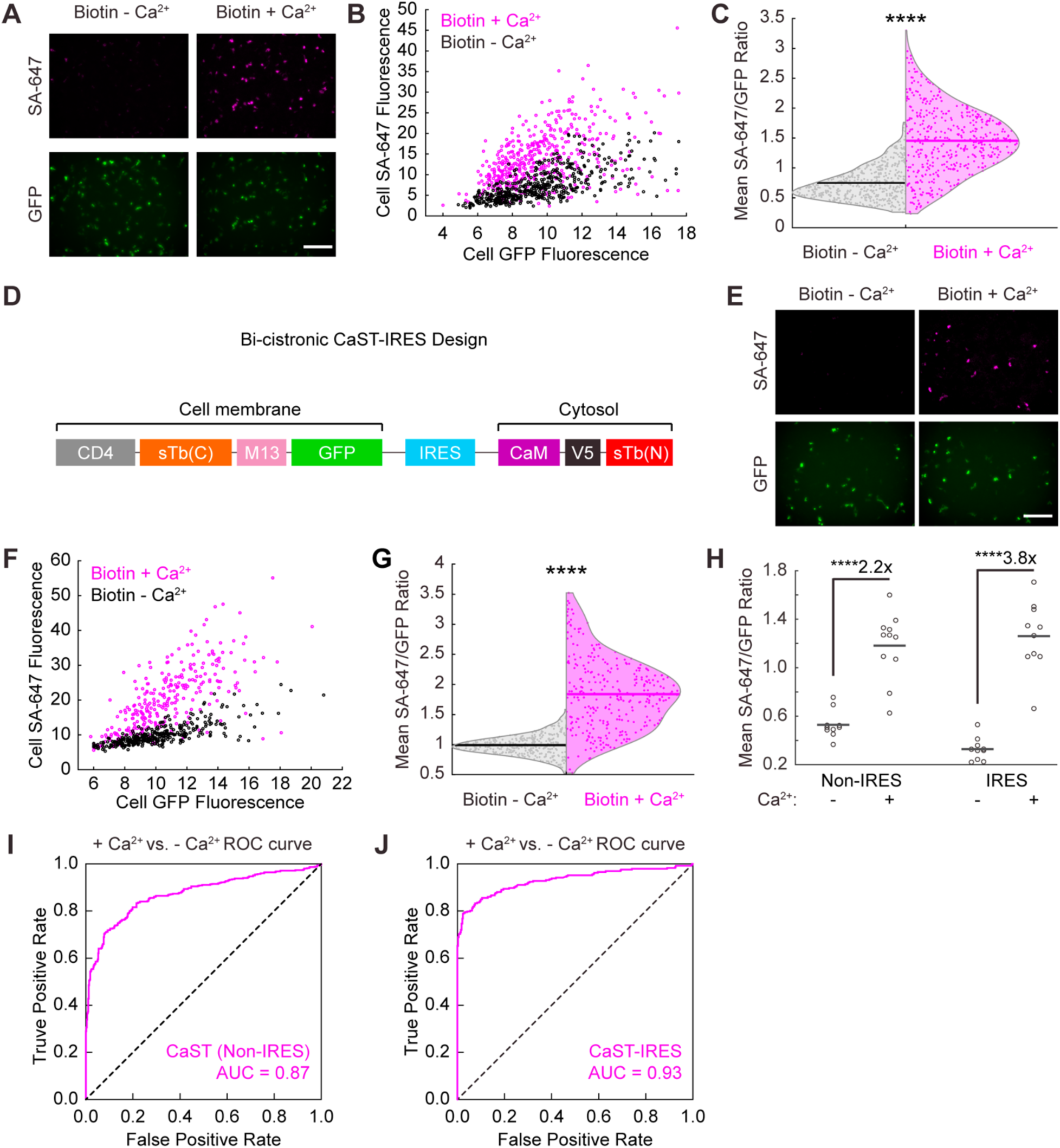
Quantification of CaST’s performance. **A)** Example images of HEK cells transfected with CaST and treated with 50 µM biotin ± Ca^2+^ (5 mM CaCl2 and 1 µM ionomycin) for 30 minutes. Top row: SA-647 staining of biotinylated proteins. Bottom row: CD4-sTb(C)-M13-GFP. **B)** Scatter plot of the mean SA-647 versus mean GFP fluorescence calculated for every GFP+ cell detected across 11 FOVs treated with either biotin + Ca^2+^ (*n* = 451 cells; Two-tailed **Pearson’s *R* = 0.50, *P* = 6.2e-30**) or biotin - Ca^2+^ (*n* = 473 cells; Two-tailed **Pearson’s *R* = 0.69, *P* = 2.6e-67**). **C)** Violin plots showing the distributions of the mean SA-647/GFP fluorescence ratio per cell from data in panel **B** (*P* = 2.0e-85, U = 27234, two-tailed Mann-Whitney U test). **D)** Schematic of the bi-cistronic CaST-IRES construct design. **E)** Example images of HEK cells transfected with CaST-IRES and treated with biotin ± Ca^2+^ for 30 minutes, as in panel **A**. **F)** Scatter plot of the mean SA-647 versus mean GFP fluorescence calculated for each GFP+ cell detected across 10 FOVs treated with either biotin + Ca^2+^ (*n* = 293 cells; Two-tailed **Pearson’s *R* = 0.72, *P*** = 1.5e-47) or biotin - Ca^2+^ (*n* = 332 cells; Two-tailed **Pearson’s *R* = 0.78, *P*** = 3.7e-69). **G)** Violin plots showing the distributions of the mean SA-647/GFP fluorescence ratio per cell from data in panel **F** (*P* = 3.1e-77, U = 6732, two-tailed Mann-Whitney U test). **H)** The FOV averages of the SA-647/GFP fluorescence ratio per cell from the non-IRES data shown in panel **B** (*n* = 11 FOVs per condition; *P* = 1.1e-5, U = 2, two-tailed Mann-Whitney U test), and the IRES data shown in panel **F** (*n* = 10 FOVs per condition; *P* = 1.1e-5, U = 0, two- tailed Mann-Whitney U test). **I,J)** ROC curves for distinguishing Ca^2+^-treated vs. non-treated cell populations based on CaST Non-IRES cells from Panel **C** (**I**; AUC = 0.87, *P* = 2.0e-85, Wilson/Brown’s method) and CaST- IRES transfected cells from Panel **G** (**J**; AUC = 0.93, *P* = 3.1e-77, Wilson/Brown’s method). All scale bars, 300 µm. \*\*\*\**P*<0.0001.

To optimize CaST delivery, we concatenated its two fragments into a bi-cistronic vector containing either a porcine teschovirus 2A peptide (P2A) coding sequence or internal ribosome entry sites (IRES). P2A and IRES are well-established strategies for co-expressing multiple proteins from a single promoter^21, 22^, ensuring that each cell expresses both fragments of CaST. We found that both CaST-P2A and CaST-IRES exhibited higher SA-647/GFP labeling with biotin and Ca^2+^ compared to with biotin alone; but the IRES version resulted in a higher biotin ± Ca^2+^ SBR (5-fold, compared to 2.7-fold; **Supplementary** Fig. 2). The IRES motif is reported to lower the expression level of the second component relative to the first^23^; this may explain why this strategy performs better than P2A, given our data showing an optimal transfection ratio of 5:2 for the two separate components of CaST (**Extended Data** Fig. 1A). For these reasons, we chose to further characterize the CaST-IRES version (**Fig. 2D-G**).

Western blot analysis confirmed that CaST-IRES cells exhibited elevated biotinylation signal compared to negative control conditions (**Extended Data** Fig. 2E). In addition, immunofluorescence analysis showed that in comparison to the non-IRES version, CaST-IRES resulted in a larger calcium-dependent fold-change in both the mean SA-647/GFP cell ratio and the mean SA-647 cell fluorescence (likely due to more controlled protein expression levels of the two components; **Fig. 2H** and **Extended Data** Fig. 3A,B). We also performed receiver operating characteristic (ROC) analyses of the SA-647/GFP ratios to evaluate CaST’s ability to discriminate between individual Ca^2+^-treated and non-treated cells. We determined an area under the curve (AUC) of 0.87 for non-IRES CaST, and an AUC of 0.93 for CaST-IRES, indicating that both versions can robustly distinguish activated versus non-activated cells (**Fig. 2I,J**).

### Temporal resolution, Ca^2+^-sensitivity, and time integration of CaST

One important requisite of our design is that both the Ca^2+^-sensing and split-TurboID reconstitution be reversible. This is so that enzyme activated during high Ca^2+^ prior to the desired biotin labeling window can split back into inactive fragments once the cytosolic Ca^2+^ returns to resting levels. To test for the reversibility of CaST, we treated HEK cells expressing CaST-IRES with Ca^2+^ for 30 minutes, washed the cells over the course of 10 minutes, and then delivered biotin for 30 minutes. We directly compared this condition to CaST-IRES cells treated with biotin alone, or with biotin and Ca^2+^. Cells treated with biotin after removal of Ca^2+^ exhibited no biotinylation, similar to the negative control (**Fig. 3A,B**). This demonstrates that CaST has a temporal resolution for detecting intracellular Ca^2+^ on the order of 10 minutes (meaning it can ignore Ca^2+^ events that occur 10 minutes prior to the desired biotin labeling window).

**Figure 3.**
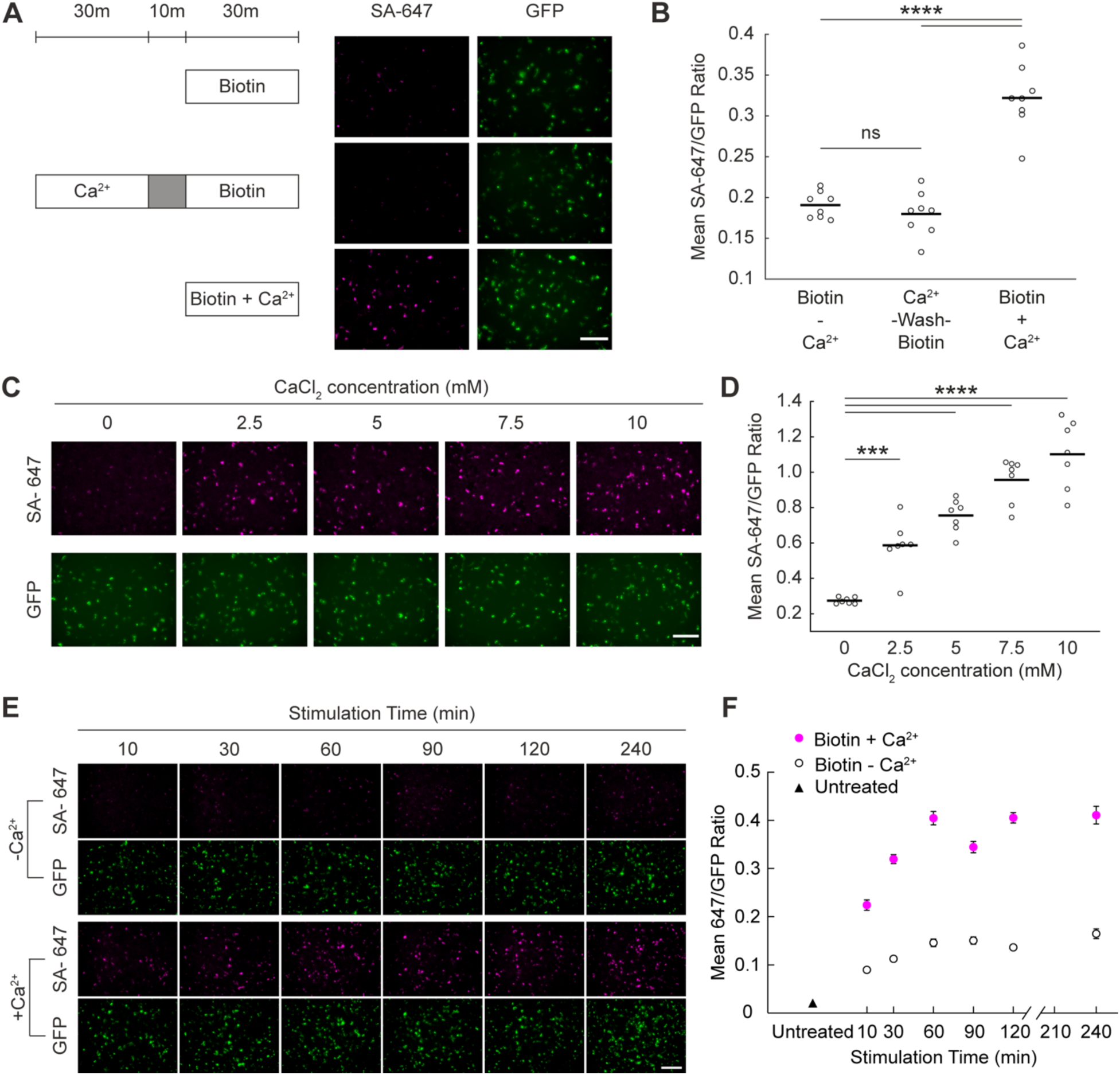
Characterization of reversibility, Ca^2+^ sensitivity, and time integration of CaST. A) To test the split enzyme’s reversibility, HEK cells were transfected with CaST-IRES and treated either with biotin alone for 30 minutes (top), with Ca^2+^ for 30 minutes followed by a 10- minute wash and then biotin for 30 minutes (middle), or with biotin + Ca^2+^ simultaneously for 30 minutes (bottom). Example images are shown for all 3 conditions with SA-647 staining of biotin and GFP expression of CaST-IRES. Scale bar, 300 µm. B) The FOV averages of the SA-647/GFP fluorescence ratio per cell for the 3 conditions shown in panel **A** (*n* = 8 FOVs per condition; Biotin - Ca^2+^ vs. Ca^2+^/wash/Biotin: *P* = 0.75; Biotin - Ca^2+^ vs. Biotin + Ca^2+^: *P* = 4.9e-8; Ca^2+^/wash/Biotin vs. Biotin + Ca^2+^: *P* = 1.3e-8; Tukey’s post-hoc multiple comparison’s test following a 1-way ANOVA, F2,21 = 56.37, *P* = 3.6e-9). C) Example FOVs of HEK cells transfected with CaST-IRES and treated with biotin and increasing concentrations of CaCl2 (and 1µM ionomycin). D) The FOV averages of the SA-647/GFP fluorescence ratio per cell for the CaCl2 concentrations shown in panel **C** (*n* = 7 FOVs per condition; 0 mM versus 2.5 mM: *P* = 7.6e-4; 0 mM versus 5 mM: *P* = 8.8e-7; 0 mM versus 7.5 mM: *P* = 5.5e-10; 0 mM versus 10 mM: *P* = 5.4e-12; Tukey’s post-hoc multiple comparison’s test following a 1-way ANOVA, F4,30 = 44.07, *P* = 3.8e-12). The FOV average SA-647/GFP ratios were linearly correlated with CaCl2 concentration (Two-tailed Pearson’s correlation coefficient *R* = 0.99, *P* = 0.001). E) Example FOVs of HEK cells transfected with CaST-IRES and treated with 50 µM biotin and ± Ca^2+^ (5 mM CaCl2 and 1 µM ionomycin) for different durations. F) The mean FOV averages of the SA-647/GFP fluorescence ratio per cell for the different stimulation times shown in panel **E** (*n* = 10 FOVs per condition). The untreated condition is shown on the left. Data is plotted as mean ± s.e.m. All scale bars, 300 µm. \*\*\**P*<0.001, \*\*\*\**P*<0.0001, ns, not significant.

Next, we asked whether the CaST signal is correlated to the levels of intracellular Ca^2+^ present in cells. We conducted a Ca^2+^ titration experiment in which we treated cells with increasing concentrations of CaCl2 in the media (along with 1 µM ionomycin and biotin) for 30 minutes. Our results demonstrated a monotonically increasing SA-647/GFP signal from 2.5 mM to 10 mM CaCl2, with a linear correlation to Ca^2+^ concentration over this range (**Fig. 3C,D**).

We also conducted temporal integration experiments to evaluate the labeling of CaST over different time exposures to a fixed Ca^2+^ concentration. CaST-IRES cells were treated with biotin ± Ca^2+^ for increasing durations of time. We found that a 10-minute stimulation period is sufficient to induce elevated SA-647/GFP labeling over background (**Fig. 3E,F**). The amount of labeling increased with longer stimulation times, saturating around 1 hour (**Extended Data** Fig. 3C,D). These results show that CaST acts as an integrator of Ca^2+^, both in terms of detecting increasing Ca^2+^ concentration, and increasing duration of Ca^2+^ exposure.

### Direct comparison of CaST to existing technologies

Compared to existing Ca^2+^-dependent integrators, CaST has two major advantages: it is non-invasive (using biotin instead of light), and it acts on rapid timescales (tags cells within minutes rather than hours). We directly compared the performance of CaST against an existing light- and Ca^2+^-dependent integrator, FLiCRE^6^. FLiCRE works via a protease-mediate release of a non-native transcription factor (e.g., Gal4) in the presence of blue light and Ca^2+^. This released transcription factor then enters the nucleus to drive expression of a modular reporter gene, such as a fluorescent protein (**Extended Data** Fig. 4A). While it has fast labeling kinetics (it can detect 30-s of blue light and elevated Ca^2+^), its read-out kinetics are slow, requiring transcription and translation of the reporter gene that can take hours to accumulate. Here we transfected HEK cells overnight with either CaST-IRES or FLiCRE and treated cells for 15 minutes with either biotin ± Ca^2+^, or light ± Ca^2+^, respectively. We then fixed cells either immediately, 2, 4, 6, or 8 hours after the stimulation, and imaged the reporter expression in each case (SA-647 for CaST, and UAS::mCherry for FLiCRE; **Fig. 4A,B**, and **Extended Data** Fig. 4B,C). Across all time points measured, we observed an increase in CaST SA-647 labeling in conditions treated with biotin and Ca^2+^ compared to with biotin alone; however, an increase in FLiCRE UAS::mCherry reporter expression was only apparent starting 6 hours after the light and Ca^2+^ stimulation (**Fig. 4C,D**). The biotin ± Ca^2+^ and light ± Ca^2+^ SBRs were calculated by normalizing the SA-647 or UAS::mCherry expression against an expression marker for each respective tool (GFP; **Fig. 4E,F**, and **Extended Data** Fig. 4D-G). Note that for the cells treated with biotin and Ca^2+^, the decrease in the SA-647/GFP ratio over the 8 hours is due to both a decrease in SA- 647 signal (protein turnover), and an increase in CaST GFP expression (**Extended Data** Fig. 4D,E).

**Figure 4.**
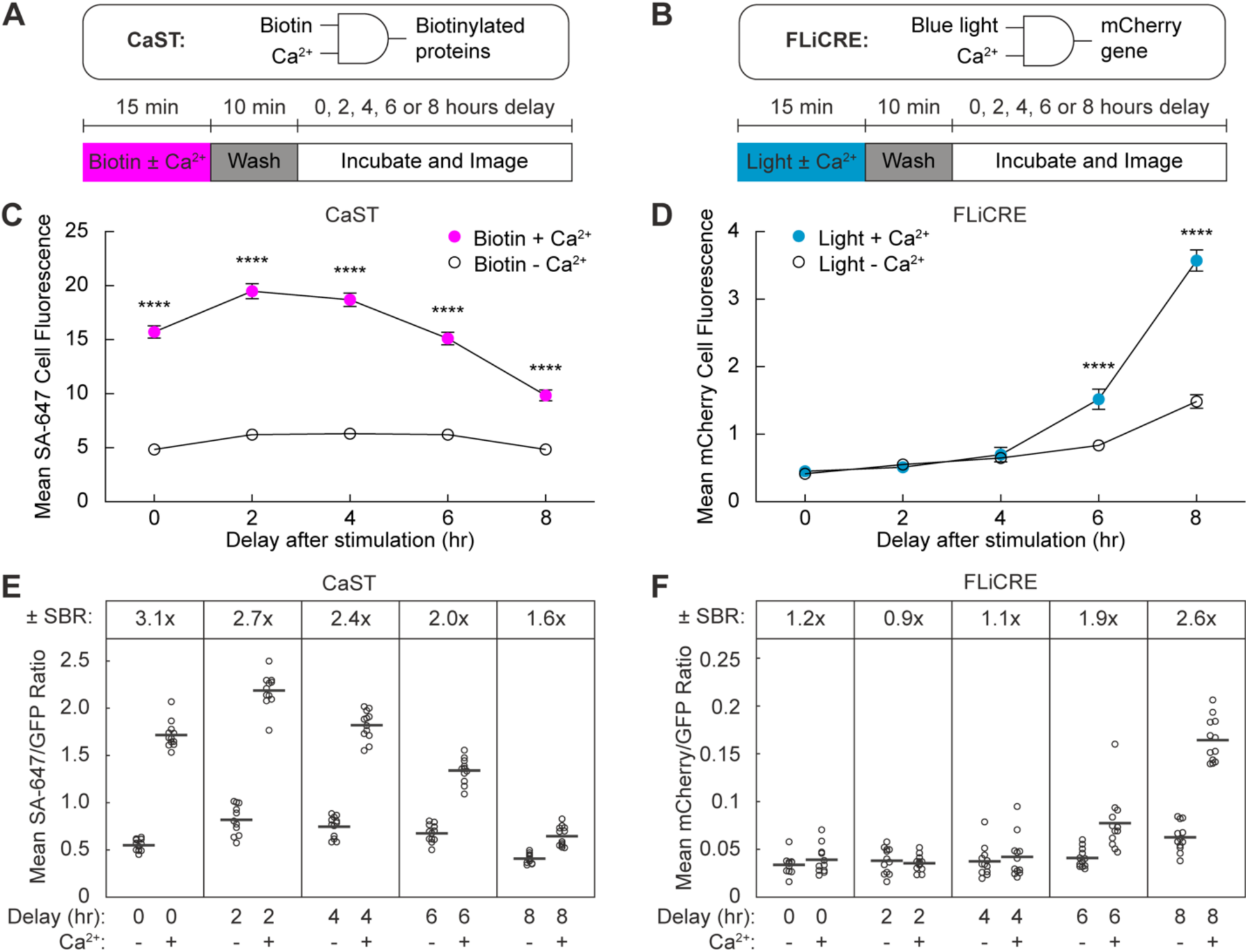
Comparison of CaST to an existing technology, FLiCRE. **A,B)** Schematics of CaST (**A**) and FLiCRE (**B**) as AND logic gates, and the experimental paradigms used to test the time course of labeling detection after biotin + Ca^2+^ (CaST) and light + Ca^2+^ (FLiCRE) stimulation. Cells were transfected with either CaST-IRES or FLiCRE (**Extended Data** Fig. 4A) components. **C)** For CaST, the FOV average of the SA-647 cell fluorescence was calculated following a variable delay period after biotin ± Ca^2+^ stimulation (*n* = 12 FOVs for conditions with 0, 4, 6, 8 hr delay; *n* = 11 FOVs for conditions with 2 hr delay; 0 hr: *P* = 1.0**e-30**; 2 hr: *P* = 3.0e-36; 4 hr: *P* = 2.4e-35; 6 hr: *P* = 3.0e-24; 8 hr: *P* = 1.4e-10; Šídák’s post-hoc multiple comparison’s test following a 2-way ANOVA, F4,108 = 25.94, *P* = 4.5e-15). **D)** For FLiCRE, the FOV average of the UAS::mCherry cell fluorescence was calculated following a variable delay period after light ± Ca^2+^ stimulation. (*n* = 11 FOVs for conditions with 0 hr delay; *n* = 12 FOVs for conditions with 2, 4, 6, 8 hr delay; 6 hr: *P* = 4.6e-5; 8 hr: *P* = 2.4e-28; Šídák’s post-hoc multiple comparison’s test following a 2-way ANOVA, F4,108 = 46.46, *P* = 1.2e- 22). **E, F)** The ± Ca^2+^ SBR of normalized reporter expression is shown for both CaST (panel **E**) and FLiCRE (panel **F**). For CaST, the SA-647 fluorescence is divided by the GFP fluorescence (*n* = 12 FOVs for conditions with 0, 4, 6, 8 hr delay; *n* = 11 FOVs for conditions with 2 hr delay). For FLiCRE, the UAS::mCherry fluorescence is divided by the GFP fluorescence (*n* = 11 FOVs for conditions with 0 hr delay; *n* = 12 FOVs for conditions with 2, 4, 6, 8 hr delay). Data are plotted as mean ± s.e.m. in **C**-**D**. \*\*\*\**P*<0.0001.

To ensure that the optimal labeling parameters were used to evaluate FLiCRE, we also stimulated cells at higher expression levels of each tool (48 hours post-transfection). We observed an increase in non-specific background labeling for both CaST-IRES and FLiCRE, even immediately post-stimulation (**Supplementary** Fig. 3A,B). Nonetheless, we were still able to observe a difference between biotin ± Ca^2+^ groups immediately after stimulation with CaST- IRES (**Supplementary** Fig. 3C-E). In contrast, FLiCRE showed no differences between light ± Ca^2+^ groups, even at 8- and 24-hours post-stimulation, due to non-specific background activation caused by high expression levels (**Supplementary** Fig. 3F-H). Previous studies have also reported that the performance of FLiCRE^6^ and related protease-dependent tools^24^ can suffer at high expression levels. These findings demonstrate both the immediacy of labeling, and robustness at various expression levels, of CaST compared to protease-driven transcriptional tools such as FLiCRE.

### Application of CaST in cultured neurons

We then asked whether CaST could detect elevated intracellular Ca^2+^ in cultured rat hippocampal neurons. We expressed the two-component version of CaST using adeno-associated viruses (AAVs) to obtain the maximal expression levels, and stimulated neurons using potassium chloride (KCl) in the presence of biotin for 30 minutes. Immunofluorescence imaging confirmed that CaST robustly tagged activated neurons treated with biotin and KCl, compared to negative control conditions (**Fig. 5A,B** and **Extended Data** Fig. 5A). Stimulation for only 10 minutes was also sufficient to drive elevated CaST labeling (**Fig. 5C,D**). Quantitative analysis of individual GFP+ neurons expressing CaST showed that ∼35% of GFP+ neurons exhibited strong SA-647 labeling in the 10-minute biotin and KCl condition (thresholded as described in **Extended Data** Fig. 5B), compared to ∼10% labeled in the biotin alone condition (white bars in **Fig. 5E**). In neurons stimulated for 30 minutes, ∼65% of GFP+ neurons exhibited strong SA-647 labeling, compared to ∼10% labeled in the biotin alone condition (grey bars in **Fig. 5E**, and **Extended Data** Fig. 5B,C). ROC analysis showed that under the 30-minute condition, CaST could distinguish KCl-stimulated versus unstimulated neurons with an AUC of 0.91 (**Extended Data** Fig. 5D). DRAQ7 staining showed no apparent cytotoxicity prior to stimulation in neurons transduced with CaST (**Extended Data** Fig. 5E,F).

**Figure 5.**
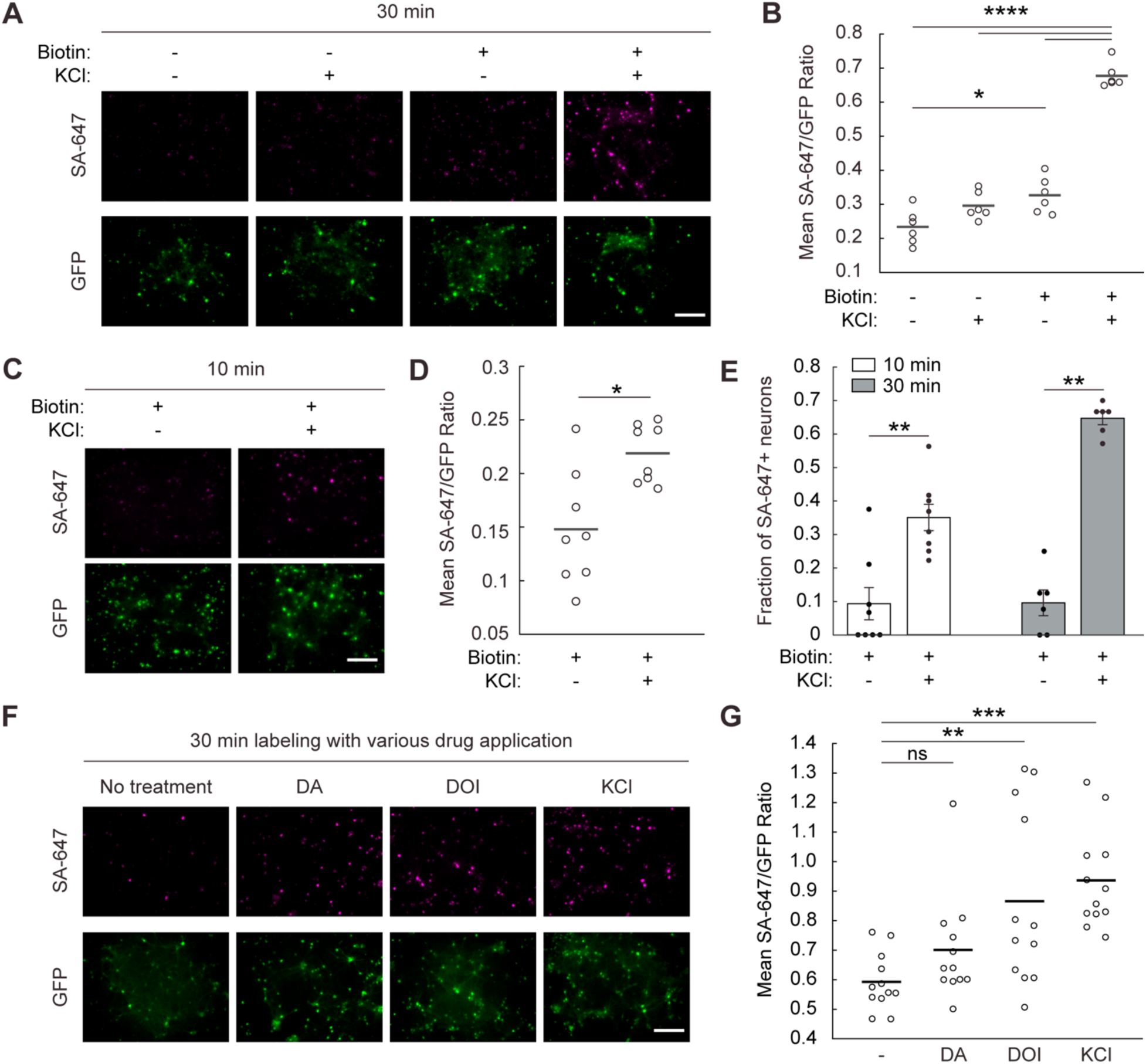
CaST performance in cultured neurons. **A)** Example FOVs of cultured rat hippocampal neurons infected with AAV2/1-Synapsin-CD4- sTb(C)-M13-GFP and AAV2/1-Synapsin-CaM-sTb(N) viruses and stimulated with ± biotin and ± KCl for 30 minutes. **B)** The FOV averages of the SA-647/GFP fluorescence ratio per cell for the data shown in panel **A** (*n* = 6 FOVs per condition; -Biotin -KCl vs. +Biotin +KCl: *P* = 1.8e-12; -Biotin +KCl vs. +Biotin +KCl: *P* = 3.0**e-11**; +Biotin -KCl vs. +Biotin +KCl: *P* = 1.4e-10; -Biotin -KCl vs. +Biotin -KCl: *P* = 0.013; Šídák’s post-hoc multiple comparison’s test following a 2-way ANOVA, F1,20 = 59.43, *P* = 2.1e-7). **C)** Example FOVs of rat hippocampal neurons infected with CaST as in panel **A** but stimulated with biotin ± KCl for only 10 minutes. **D)** The FOV averages of the SA-647/GFP fluorescence ratio per cell for the data shown in panel **C** (*n* = 8 FOVs per condition; *P* = 0.015, U = 9, two-tailed Mann-Whitney U test). **E)** Fraction of all GFP+ neurons that are also SA-647+ (defined as having an SA-647 fluorescence value greater than the 90^th^ percentile of neurons in the biotin - KCl group). Data is quantified for the 10-minute labeling experiment shown in panel **C** (*n* = 8 FOVs per condition; *P* = 0.003, U = 5, two-tailed Mann-Whitney U test) and for a replicated 30-minute labeling experiment shown in **Extended Data** Fig. 5B,C (*n* = 6 FOVs per condition; *P* = 0.002, U = 0, two-tailed Mann-Whitney U test). Data is plotted as mean ± s.e.m. **F)** Example FOVs of rat hippocampal neurons infected with CaST as in panel **A** and treated with 50 µM biotin and 10 µM dopamine (DA), 10 µM DOI, or 30 mM KCl for 30 minutes. **G)** The FOV averages of the SA-647/GFP fluorescence ratio per cell for the conditions shown in panel **F** (*n* = 12 FOVs per condition; -KCl vs. DA: *P* = 0.547; -KCl vs. DOI: *P* = 0.008; -KCl vs. KCl: *P* = 6.5e-4, Tukey’s post-hoc multiple comparison’s test following a 1-way ANOVA, F3,44 = 7.373, *P* = 4.2e-4). All scale bars, 300 µm. \**P*<0.05, \**\*P*<0.01, \*\*\**P*<0.001, \*\*\*\**P*<0.0001, ns, not significant.

Next, we explored the relationship between intracellular Ca^2+^ changes and CaST labeling using the real-time fluorescent calcium indicator RCaMP2^4^. We co-infected RCaMP2 and CaST AAVs in neurons, and mildly stimulated them using a media change. We found that neurons exhibiting greater Ca^2+^ changes in response to stimulation (quantified by the RCaMP2 mean dF/F peak height) also displayed stronger SA-647/GFP CaST labeling (**Extended Data** Fig. 6). This demonstrates on a cell-by-cell basis that CaST labeling is correlated to intracellular calcium level changes. To additionally demonstrate the specificity of CaST labeling, we used a red-shifted optogenetic cation channel, bReaChES^25^, to achieve spatially targeted neuron stimulation using orange light. We selectively stimulated neurons within a narrow ∼1mm width slit through the bottom of the culture dish. We observed an increased SA-647/GFP cell ratio only within the subregion of the FOV exposed to orange light, while KCl stimulation resulted in uniform CaST biotinylation across the dish (**Extended Data** Fig. 7).

Lastly, we investigated the potential for CaST to detect elevated intracellular Ca^2+^ in response to physiological stimuli, such as exposure to neuromodulators or pharmacological agents. We treated neurons with biotin and either 10 µM dopamine (DA), or 10 µM 2,5-dimethoxy-4- iodoamphetamine (DOI; a serotonin 5-HT type 2A/C receptor agonist), for 30 minutes. Both dopamine receptors and 5-HT2A receptors are known to be expressed in rat hippocampal neurons^26–28^. Dopamine receptors are either Gi or Gs protein-coupled receptors, while 5-HT2A receptors are Gq protein-coupled receptors^29, 30^. Whereas activation of Gi/Gs protein signaling primarily modulate cAMP with variable effects on intracellular Ca^2+^ levels, Gq protein signaling should directly elevate intracellular Ca^2+^ levels^31, 32^. We found that dopamine treatment did not increase SA-647 labeling relative to a biotin alone vehicle control; however, biotin and DOI drove an increase in SA-647 labeling (**Fig. 5F,G**). RCaMP2 imaging in neurons treated under experimentally matched conditions confirmed that both DOI and KCl (but not DA) induce an increase in Ca^2+^ activity (**Extended Data** Fig. 8). These results demonstrate that CaST can be applied to reveal differential responses to pharmacological compounds among a population of neurons containing multiple neuronal subtypes and expressing different receptors.

### Identifying psilocybin-activated neurons during head-twitch

We next applied CaST in vivo to record cellular activity in behavioral contexts that have previously been challenging to record from with existing methods. Psychedelics are an emerging class of therapeutics that are known to promote neuroplasticity in the prefrontal cortex^33–35^ and produce positive behavioral adaptations in animal models of neuropsychiatric disorders^36–39^. These compounds also drive hallucinations in humans; therefore a major pressing area of research is to understand whether the hallucinogenic and therapeutic aspects of psychoplastogens can be decoupled^40^. The main hallucinogenic behavioral correlate of psychedelic drugs in animal models is the head-twitch response (HTR)^41, 42^ – a rhythmic rotational head movement that is tightly coupled to 5-HT2A receptor activation and correlated with hallucinogenic potential in humans^42^. Critically, the HTR measurement requires free movement of the animal’s head, precluding its measurement in head-fixed rodents under a microscope during cellular-resolution neuronal recordings. We posited that CaST could be applied to directly correlate cellular neuronal activity with the psychedelic-induced HTR in vivo.

We used CaST to measure how psilocybin, a potent and therapeutically relevant psychedelic, modulates population activity in the medial prefrontal cortex (mPFC) of untethered mice during the simultaneous measurement of the HTR. Notably, there have been conflicting reports as to whether 5-HT2A receptor agonists increase^43–45^ or decrease^46, 47^ population neuronal activity in the cortex. This could be in part due to the different sensitivities in the previous recording methods used.

We expressed CaST viruses under a pan-neuronal synapsin promoter in the mPFC to identify what percentage of neurons are activated during acute psilocybin injection. Mice expressing CaST were injected with a single intraperitoneal (IP) injection of biotin and saline, or biotin and psilocybin. Mice were video recorded to quantify the number of head-twitches displayed following the drug treatment. 1 hour later, mice were sacrificed, and mPFC slices were stained for SA-647 (**Fig. 6A**). We first asked how psilocybin modulated mPFC neuronal activity using CaST as the read-out. Immunohistochemistry showed an increase in SA-647 labeling in mice treated with psilocybin, compared to control mice (**Fig. 6B**). Image quantification showed that individual CaST-expressing GFP+ neurons in psilocybin-treated mice exhibited increased SA- 647 fluorescence compared to neurons in saline-treated mice (**Fig. 6C**). The normalized SA- 647/GFP cell ratio averaged across FOVs was also elevated in psilocybin- versus saline-treated mice (**Fig. 6D**). CaST identified that ∼70% of GFP+ neurons in the mPFC exhibited strong SA- 647 labeling following psilocybin treatment (**Fig. 6E**). Mice not expressing CaST, but injected with biotin and psilocybin, did not exhibit SA-647 labeling (**Extended Data** Fig. 9A,B).

**Figure 6.**
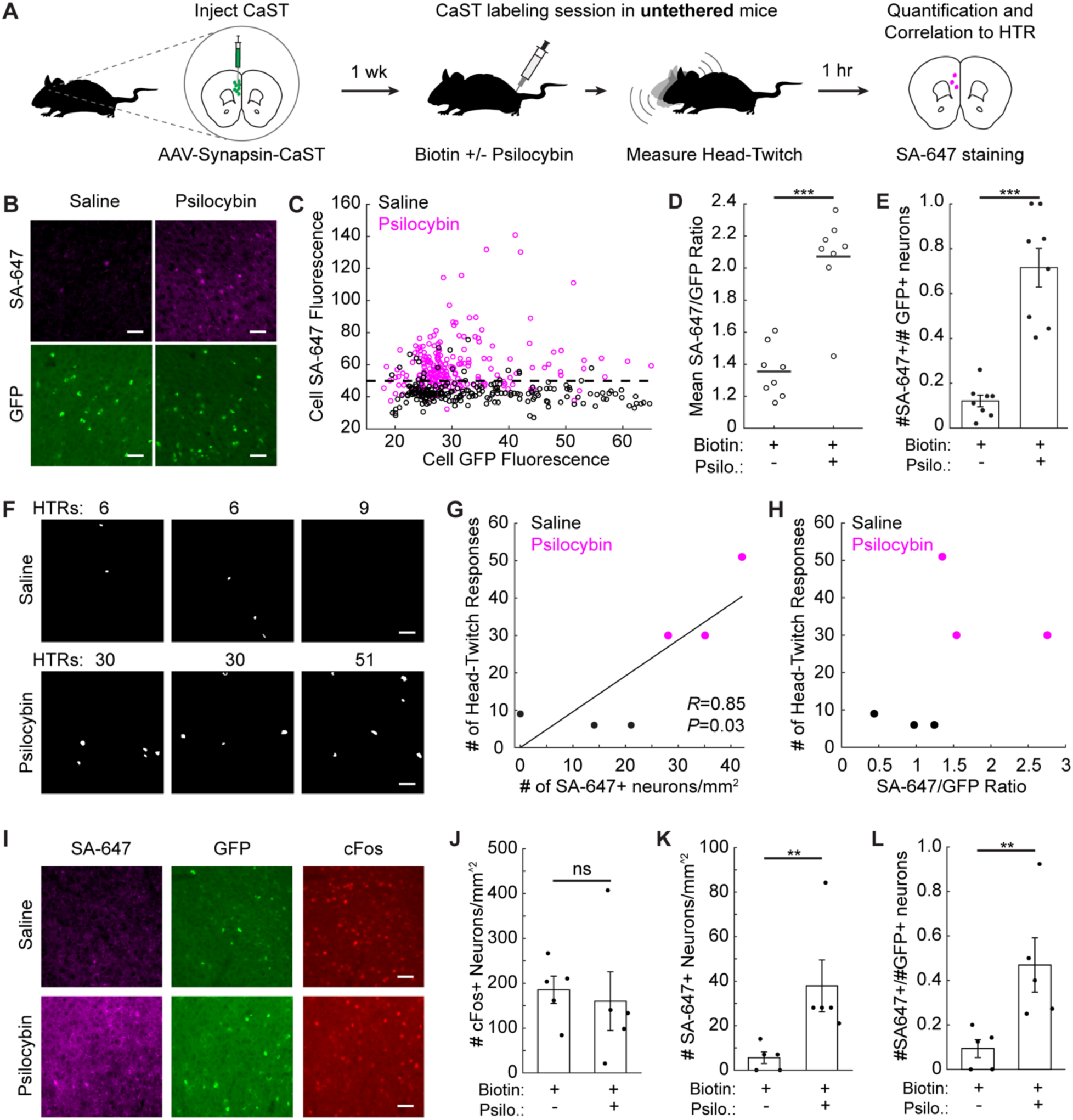
Non-invasive identification of psilocybin-activated neurons in vivo. **A)** Schematic for using CaST to tag psilocybin-activated neurons during HTR measurement. **B)** Example mPFC images of SA-647 and CaST GFP fluorescence, for mice injected with biotin+saline, or biotin+psilocybin. **C)** Mean SA-647 vs. GFP fluorescence for each GFP+ neuron detected in biotin+saline mice (*n* = 218 neurons from 3 mice) or biotin+saline injected mice (*n* = 220 neurons from 3 mice). The horizontal dashed line indicates the 90^th^ percentile threshold value of all SA-647 neurons in the biotin+saline group. **D)** FOV averages of the SA-647/GFP fluorescence ratios from panel **C** (*n* = 8 FOVs pooled from 3 mice in both conditions; *P* = 6.2e-4, U = 38, two-tailed Mann-Whitney U test). **E)** Fraction of all GFP+ neurons that are SA-647+ (thresholded using the dotted line in panel **C**; *n* = 8 FOVs pooled from 3 mice in both conditions; *P* = 1.6e-4, U *=* 36, two-tailed Mann-Whitney U test). **F)** Cell masks of SA-647+ mPFC neurons identified during HTR measurements. FOVs with the same number of HTRs were taken from the same mice, but from independent CaST injections on opposite hemispheres. **G)** Number of HTRs vs. the number of SA-647+ neurons/mm^2^ for data shown in panel **F** (*n* = 6 FOVs from independent CaST injections; Two-tailed Pearson’s correlation coefficient *R* = 0.85, *P* = 0.03). **H)** Number of HTRs vs. the mean cell SA-647/GFP ratio for data shown in panel **F**. **I)** Example mPFC images of CaST GFP, SA-647 staining, and c-Fos staining in mice treated with biotin+saline, or biotin+psilocybin. **J-L)** Number of c-Fos+ neurons/mm^2^ (**J**), SA-647+ neurons/mm^2^ (**K**), or SA-647+ divided by GFP+ neurons/mm^2^ per FOV (**L**), in biotin+saline vs. biotin+psilocybin injected mice (*n* = 5 mice per condition; (**J**) *P* = 0.42, U = 8, two-tailed Mann-Whitney U test; (**K**) *P* = 0.0079, U *=* 0, two-tailed Mann-Whitney U test; (**L**) *P* = 0.0079, U *=* 0, two-tailed Mann-Whitney U test). All scale bars, 50 µm. Data are plotted as mean ± s.e.m. in **E**, **J-L**. \*\**P*<0.01, \*\*\**P*<0.001, ns, not significant.

No prior reports have measured Ca^2+^ activity in the mPFC following psilocybin. Thus, to validate our finding that a large fraction of mPFC neurons were activated by psilocybin using CaST, we performed two-photon (2P) scanning microscopy in head-fixed mice. We injected mice in the mPFC with an AAV encoding the Ca^2+^ indicator GCaMP6f and implanted a 1.0 mm diameter Gradient-Index (GRIN) lens to optically access neurons ∼2.5 mm deep in the brain (**Extended Data** Fig. 9C). 2P imaging was conducted in mice immediately after administering saline or psilocybin (**Extended Data** Fig. 9D-F). Similar to our findings with CaST, 2P imaging identified a substantial population of neurons in mPFC activated by psilocybin (**Extended Data** Fig. 9G,H).

While the 2P GRIN lens imaging validated our CaST findings, this methodology inflicts damage to the surrounding tissue, and it also requires head-fixation to record activity. It is impossible to measure the HTR during head-fixed 2P imaging, and this behavior may also be hampered using miniaturized head-mounted microscopes^48^. Indeed, the HTR has thus far only been quantified during bulk fiber photometry recordings^49^, which lacks cellular resolution. Here we were able to simultaneously record the HTR during CaST labeling in mice injected with either saline or psilocybin. We then measured the amount of neuronal activity induced in the mPFC as a function of the HTR observed (**Fig. 6F**). The number of SA-647+ neurons was positively correlated with the number of HTRs measured during the recordings (**Fig. 6G**). The mean SA- 647/GFP fluorescence ratio of all neurons was also higher in mice exhibiting psilocybin-induced HTRs (**Fig. 6H**).

To compare these results to the current best-in-class, activity-dependent antibody staining method compatible with untethered mice, we repeated our CaST experiment while staining for cFos. We found that cFos staining did not show a relative increase in labeling in the mPFC following psilocybin injection, due to high background staining in this region even with a saline injection (**Fig. 6I-J** and **Extended Data** Fig. 10A-D). As previously reported^45^, cFos staining showed increased labeling in the somatosensory cortex (SSC) of psilocybin-treated mice (**Extended Data** Fig. 10E,F). This result confirms that there were no issues with our cFos staining protocol and suggests that a major limitation of cFos is its variability across brain regions. In contrast, CaST showed an increased number of SA-647+ neurons in mice treated with psilocybin compared to saline control, even when normalizing for the total number of CaST- expressing GFP+ cells in each FOV (**Fig. 6K,L**). All together, these results establish CaST as a sensitive technology for rapid and non-invasive tagging of activated neurons in untethered mice – providing a powerful and complementary strategy to existing molecular tools for recording and identifying activated neurons in vivo.

## DISCUSSION

Here we demonstrate an enzymatic approach for stably tagging activated cells with a bio- compatible handle both in vitro and in vivo. Our method, CaST, acts as a Ca^2+^-dependent ligase, labeling itself and nearby proteins with a biotin tag in living cells. Endogenous biotin levels in cells and in the brain are low enough that CaST requires exogenously delivered biotin to robustly tag proteins. Thus, CaST acts as a time-gated integrator of Ca^2+^ concentration, driving the accumulation of biotinylated proteins during a specified labeling period. Importantly, CaST reconstitution is reversible, and following a period of Ca^2+^-activation, it can be reset to its inactivated state within 10 minutes. Due to its tight temporal resolution and Ca^2+^ sensitivity, CaST could robustly detect mPFC neurons activated by psilocybin in vivo during the simultaneous measurement of the HTR (which could not be detected using cFos labeling).

CaST reported that a large fraction of neurons in the prelimbic mPFC are activated by psilocybin in mice. This was corroborated by our own secondary validation studies using single-cell Ca^2+^ imaging. A previous study in rats using in vivo electrophysiology reported that another 5-HT2A receptor agonist, DOI, primarily inhibits mPFC population activity in rats^46^. It is possible that in vivo electrophysiology recordings may be biased towards more highly active neurons at baseline (missing more silent neurons that are activated by psychedelics); or that DOI and psilocybin have different effects on population activity in mPFC. Another recent study using cFos labeling following psilocybin injection (1 mg/kg) reported that they could detect elevated cFos labeling in the mPFC only when using one of the two different imaging modalities that they tested^45^; in addition, our own studies here showed that we could not detect a difference in mPFC cFos labeling following psilocybin injection. This suggests that the neuronal activity changes driven by psilocybin in the mPFC are challenging to tag using IEG-based approaches and are better detected using Ca^2+^-based approaches such as CaST.

In contrast to existing light-gated methods for Ca^2+^-dependent labeling^5, 6, 8^, a major advantage of CaST is that it is essentially non-invasive, requiring only the brief injection of a biotin solution IP in mice which can rapidly cross the blood-brain barrier. This is particularly advantageous in deep tissue, where light must be delivered through fiber implantation, limiting the recording area and the range of tool usage. It could also be advantageous in other areas of the body, such as the pancreas or in the spinal cord, where it is not possible to deliver blue light. As TurboID can already be used for cell-type-specific biotin proximity labeling across the entire brain^50^, it is feasible that CaST can also be scaled as a brain-wide activity-dependent labeling tool in the future, with the use of PhP.eB^51^ viruses or transgenic lines. Although we did not observe overt cytotoxicity in neurons expressing CaST (nor has it been reported for transgenic TurboID mice^50^), additional assays of neuron physiology and cell health should be performed if expressing CaST for prolonged periods of time in the brain.

In addition, because the biotin molecule is directly attached to already-expressed proteins, this enables the immediate read-out of activated neurons after the labeling period with CaST. This contrasts with transcriptional-based reporters, including both Ca^2+^- and IEG-dependent systems, which require multiple hours before activated cells can be identified by the reporter RNA or protein. The immediacy of labeling, along with the scalability of CaST, would be particularly useful for registering cellular activity history with other spatial molecular imaging modalities, such as MERFISH^52^ or STARmap^53^ for in situ transcriptomics, or MALDI-IHC^54^ for spatial proteomics. Because CaST affords a stable read-out of neuron activation following tissue fixation, one could apply spatial omics approaches to identify acutely induced changes in gene expression or protein localization caused specifically in this subset of activated neurons. The spatial distribution of fluorescently stained biotinylated proteins could be imaged and registered to the in situ omics data, or a streptavidin-oligo bound to biotinylated proteins could be detected during in situ sequencing^55^.

However, despite the advantages of CaST, it is important to note its limitations in comparison to existing tools. For example, some users may still require the fast labeling windows afforded by light- and calcium-gated tools, to tag neurons activated during acute behaviors which cannot be isolated during the relatively broad biotin labeling period. In addition, we note that the biotin labeling is an “end point experiment” where cells must be fixed and stained for imaging or collected and lysed for protein analysis. Although biotinylated cells can be subjected to subsequent downstream molecular or chemical analysis, proximity labeling itself does not enable further manipulation or genetic access to biotin-tagged neurons. Consequently, CaST is intended to complement, rather than replace, existing tools that can activate induced transcription factors and drive the expression of proteins such as opsins for downstream neuronal manipulation. Finally, there are several considerations for using CaST in vivo, including the relative time window of tagging based on the biotin injection, the strength of the behavioral stimulus required to induce strong enough CaST labeling, and the need for negative controls animals to determine the appropriate thresholds for detecting true CaST signal (**Supplementary Note 1**).

Finally, although here we primarily highlight CaST as an enzymatic-based activity integrator, it is worth noting that it could also be used as a Ca^2+^-dependent proximity labeling by analyzing biotinylated proteins (see **Fig. 1D** and **Extended Data** Figure 2). Thus, future studies could also enrich CaST-tagged proteins for analysis with mass spectrometry to examine activity-dependent differences in subcellular protein expression among unique neuronal cell-types.

## ACKNOWLEDGEMENTS

We thank Alice Y. Ting (Stanford University) for the generous gift of plasmids used as templates in this study. We thank Jessie Muir (UC Davis) for assistance with GRIN lens surgeries for in vivo mouse imaging. We thank Hunter T. Warren (UC Davis) and Winston L. Chow (UC Davis) for synthesizing psilocybin. This work was supported by a Burroughs Wellcome Fund Career Awards at the Scientific Interface (#1019469, C.K.K.), a NARSAD Young Investigator Grant (#30238, C.K.K.) the Searle Scholars Program (#SSP-2022-107, C.K.K.), the Arnold and Mabel Beckman Foundation (Beckman Young Investigator Award, C.K.K.), the National Institutes of Health (DP2MH136588, C.K.K.; and R35GM148182, D.E.O.), and the Boone Family Foundation (D.E.O). R.Z. was supported by a National Science Foundation (NSF) Research Traineeship (NeuralStorm, #2152260). M.A. was supported by an NSF Graduate Research Fellowship Program (#000895154). J.C. was supported by a National Institutes of Health T32 Training Program (UC Davis Learning, Memory and Plasticity Training Program, T32 MH112507).

## AUTHOR CONTRIBUTIONS STATEMENT

R.Z. and C.K.K. conceived the tool design and conceptualized the project. R.Z. performed protein engineering, cultured HEK cell and neuron experiments, image analysis, and data quantification. M.A. performed and oversaw all Western blot analysis and led the design and implementation of mouse in vivo CaST experiments, with assistance from I.A.K. I.A.K. analyzed CaST head-twitch data and assisted with CaST in vivo surgeries and labeling injections. S.L. performed Western blot experiments and generated AAVs for CaST expression. J.A.C. performed GRIN lens surgeries for in vivo mouse imaging and collected 2P imaging data, with assistance from I.A.K. S.S.V. assisted with HEK cell assays characterizing the different versions of CaST. D.E.O. and I.A.K. provided psilocybin along with critical experimental design input for the in vivo GCaMP imaging and CaST labeling during psilocybin injection. R.Z. led the figure preparation with guidance from C.K.K. R.Z. and C.K.K. wrote the manuscript with contributions from all authors.

## METHODS

### Cloning

All constructs used or developed in this study are listed in **Supplementary Table 1**. For all constructs, the vectors were double digested with restriction enzymes (New England BioLabs; NEB) following the standard digest protocols. PCR fragments were amplified using Q5 polymerase (M0494S, NEB). Both vectors and PCR fragments were purified using gel electrophoresis and gel extraction (28706, Qiagen) and were ligated using Gibson assembly (E2611S, NEB). Ligated plasmids were introduced into NEB Stable Competent *E. coli* (C3040H, NEB) via heat shock following the manufacture’s transformation protocol. Plasmids were amplified using NEB 5-alpha Competent *E. coli* (C2987H, NEB) and Plasmid Miniprep Kit (27106, Qiagen).

### AlphaFold2 protein structure predictions

To generate AlphaFold2^57^ predicted structures of CaST in Figure 1, ColabFold^56^ (v1.5.5) was used using default settings. To generate the predicted structures of the two halves of CaST in isolation, each half’s amino acid sequence was entered individually, to predict each structure separately. To generate the predicted structure of the two halves of CaST in complex, the two amino acid sequences were entered, separated by a “:” to specify inter-protein chainbreaks for modeling complexes (e.g., heterodimers). Predicted structures were re-colored using Pymol (v2.5.2).

### Mammalian cell culture and transfection

Human Embryonic Kidney cell 293T (CRL-3216, ATCC; no additional verification performed) were cultured as a monolayer in Dulbecco’s Modified Eagle Medium (DMEM; D5796-500ML, Sigma-Aldrich), supplemented with 10% Fetal Bovine Serum (FBS; F1051-500ML, Sigma-Aldrich) and 1% (v/v) penicillin/streptomycin (P/S; 15070063, 5,000 U/mL, Life Technologies) (complete DMEM). Cells were cultured in 100 mm TC treated dishes (353003, Falcon) and maintained in the cell culture incubator at 37°C with humidified 5% CO2 and subcultured when they reached 80%–90% confluence using Trypsin (T2610-100ML, Sigma-Aldrich).

For immunofluorescence experiments, cells were plated in 48-well plates pre-treated with 50 mg/mL human fibronectin (FC010-10MG, Millipore) and incubated for 24 hr. Cells at ∼80% confluency were transfected with DNA plasmids using polyethyleneimine max according to the manufacturer’s manual (PEI Max; 24765-1, Polysciences). For CaST experiments, cells were transfected with 50 ng CD4-sTurboID (C)-M13-GFP, 20 ng CaM-V5-sTurboID (N), and 0.8 µL PEI Max per well. For CaST-IRES experiments, cells were transfected with 50 ng CD4- sTurboID (C)-M13-GFP-IRES-CaM-V5-sTurboID (N) and 0.8 µL PEI Max per well. Cells were incubated at 37°C overnight for 15-16 hr prior to stimulation.

For confocal experiments, cells were plated in 35 mm glass bottom dishes (D35-14-1-N, Cellvis) pre-treated with 50 mg/mL human fibronectin and incubated for 24 hr. Cells at ∼80% confluency were transfected with DNA plasmids using PEI Max according to the manufacturer’s manual. For CaST experiments, cells were transfected with 250 ng CD4-sTurboID (C)-M13-GFP, 125 ng CaM-V5-sTurboID (N), and 8 µL PEI Max per well. Cells were incubated at 37°C overnight for 15-16 hr prior to stimulation.

### HEK293T cell CaST / CaST-IRES experiments

CaST or CaST-IRES HEK cells were stimulated 15-16 hour following transfection. Stimulation master mixes were made by adding CaCl2 (21115-100ML, Sigma-Aldrich), Ionomycin (I3909- 1ML, Sigma-Aldrich), and Biotin (B4639, Sigma-Aldrich; dissolved in DMSO as 100mM stock) to complete DMEM. The stimulation mixtures were then added to the cell and the cells were incubated at 37°C for 30 min for CaST/CaST-IRES labeling. For +Ca^2+^ conditions, we used a final concentration of 5 mM Ca^2+^ and 1 µM Ionomycin. For +biotin conditions, we used a final concentration of 50 µM biotin. After incubation for the indicated time, the solution was removed in each well. Cells were washed once with warm complete DMEM and twice with warm DPBS (D8537-500ML, Sigma-Aldrich). Cells were subsequently fixed with 4% (v/v) paraformaldehyde (PFA; sc-281692, Santa Cruz Biotechnology) in DPBS at room temperature for 10 min followed by two washes with DPBS. After that, cells were permeabilized with ice-cold methanol at -20°C for 8 min followed by two washes with DPBS at room temperature. Cells were incubated with blocking solution (1% w/v BSA; BP1600-100, Fisher Scientific, in DPBS) for 45 min at room temperature; followed by a 1 hr incubation in primary antibody solution at room temperature (1:1,000 mouse anti-v5; R96025, Invitrogen). Cells were washed twice with DPBS and incubated with secondary antibody (1:1,000 donkey anti-mouse 568; ab175472, Abcam) and streptavidin-Alexa Fluor647 (1:5,000; S32357, Invitrogen) for 30 min at room temperature. Cells were then washed three times with DPBS and imaged by epifluorescence microscopy (see “***Immunofluorescence imaging***”).

For CaST variable transfection ratio experiments, cells at ∼80% confluency were transfected with different ratios of CD4-sTurboID(C)-M13-GFP and CaM-V5-sTurboID(N) using PEI Max according to the manufacturer’s manual. Single fragment CaST experiments were performed by transfecting cells with only one CaST fragment (either 50 ng CD4-sTb(C)-M13-GFP or 20 ng CaM-V5-sTb(N)) or both CaST fragments as positive control. Reversibility experiments were performed by treating the cells with 5 mM Ca^2+^ and 1 µM ionomycin without biotin at 37°C for 30 min followed by two DPBS washes to wash away Ca^2+^. Then, the cells were treated with 50 µM biotin without Ca^2+^ for another 30 min. Ca^2+^ titration experiments were performed by treating the cells with different CaCl2 concentrations ranging from 0 to 10 mM and 1 µM ionomycin at 37°C for 30 min. Time integration experiments were performed by treating the cells with 5 mM Ca^2+^ and 1 µM ionomycin with or without biotin at 37°C for diverse durations ranging from 10 to 240 minutes. Stimulation, fixation, and staining were performed as described above.

### HEK293T FLiCRE / CaST-IRES comparison experiment

Transfection of HEK293T cells with FLiCRE was performed following published protocols^6^. In detail, cells were transfected with FLiCRE constructs using PEI max (20 ng UAS-mCherry, 30 ng Calmodulin-TEVp, 50 ng CD4-MKII-LOV-TEVcs-Gal4, and 0.8 µL PEI max per well in 48- well plates) at ∼80% confluency and were immediately wrapped in foil and incubated overnight for 8-9 hr. Transfection of CaST-IRES was performed as described above. FLiCRE expressing cells were treated with continuous blue light ± Ca^2+^ (6 mM Ca^2+^ and 1 µM ionomycin) for 15 min at 37°C. CaST-IRES cells were treated with biotin ± Ca^2+^ with a final concentration of 5 mM Ca^2+^, 1 µM ionomycin, and 50 µM Biotin and incubated at 37°C for 15 min. After stimulation, cells were washed twice with warm DPBS and incubated with DPBS at 37°C for 10 min. After the washes, the DPBS was removed and replaced with 200 µL complete DMEM for further incubation. The cells were fixed either immediately, 2, 4, 6, or 8 hr after the stimulation. Staining was performed after all the cells had been fixed following the methods described above. The biotinylated proteins (SA-647, CaST-IRES) and expression of the reporter (UAS::mCherry, FLiCRE) were imaged by fluorescence microscopy.

For extended post-transfection incubation, cells were transfected with CaST-IRES or FLiCRE constructs as described above and were incubated at 37°C for 48 hrs before stimulation. Cells were stimulated, washed, and incubated as described above. The cells were fixed either immediately, 8, or 24 hrs after the stimulation. Staining was performed after all the cells had been fixed following the methods described above. The biotinylated proteins (SA-647, CaST- IRES) and expression of the reporter (UAS::mCherry, FLiCRE) were imaged by fluorescence microscopy.

### Western blot analysis of CaST

HEK293T cells expressing CaST or CaST-IRES were stimulated and labeled with biotin as described above in 6-well plates and were subsequently washed 3 times with 1 mL DPBS. Cells were then detached from the well by gently pipetting 1 mL of ice-cold DPBS onto the cells and were pelleted by centrifugation at 300 g at 4°C for 3 minutes. The resulted supernatant was removed, and the pellet was lysed on ice for 10 minutes using 100 μL RIPA lysis buffer (50 mM Tris pH 8, 150 mM NaCl, 0.1% SDS, 0.5% sodium deoxycholate, 1% Triton X-100) supplemented with protease inhibitor cocktail (P8849, Sigma-Aldrich) and 1 mM PMSF (82021, G-Biosciences). The cell lysates were clarified by centrifugation at 13,000 g at 4°C for 10 minutes. The supernatants were collected and mixed with 4x protein loading buffer (PLB). The resulted samples were boiled at 95°C for 10 minutes prior to separation on a 10% precast SDS-PAGE gel. Separated proteins on SDS-PAGE gels were transferred to a nitrocellulose membrane in transfer buffer on ice. The membrane was removed after the transfer and incubated in 5 mL Ponceau stain (0.1% (w/v) Ponceau in 5% acetic acid/water for 5 minutes. The Ponceau stain was then removed with diH2O and then rinsed with TBST (Tris-buffered saline, 0.1% Tween 20). The membrane was then blocked in 5% (w/v) nonfat dry milk in TBST at room temperature for 1 hour and washed 3 times with TBST for 5 minutes each. To detect biotinylated proteins, the membrane was incubated in 10 mL of 3% BSA/TBST (w/v) with 2 µL streptavidin-HRP (1:5,000; S911, Invitrogen) for 1 hour at room temperature. The blot was washed 3 times with TBST for 5 minutes and was developed with Clarity Max Western ECL Substrate (Bio-Rad) for 1 minute before imaging. To detect the V5 tag, the membrane was incubated in 10 mL of 3% BSA/TBST (w/v) with 1 µL mouse anti-V5 primary antibody (1:10,000; R96025, Invitrogen) for 1 hour at room temperature. The blot was washed 3 times with TBST for 5 minutes and was incubated with 10 mL of 3% BSA/TBST (w/v) with 1 µL anti-mouse-HRP secondary antibody (1:10,000; 170-6516, Bio-Rad) for 30 minutes at room temperature. The blot was washed 3 times with TBST for 5 minutes and was developed with Clarity Max Western ECL Substrate (Bio-Rad) for 1 minute before imaging. The blots were imaged on a Bio-Rad Chemi-Doc XR gel imager. The raw Western blot images were quantified using Fiji/ImageJ v2.9.0 (the exact parameters used for each blot is shown in the relevant figure legends).

### AAV1/2 virus production, concentration, and titration

The production and concentration processes for AAV viruses were conducted following a previously reported method^5, 60^. In detail, HEK293T cells at ∼80% confluency in 3 T-150 flasks were transfected with 5.2 µg AAV vector, 4.35 µg AAV1 plasmid, 4.35 µg AAV2 plasmid, 10.4 µg DF6 AAV helper plasmid, and 130 µL PEI Max solution per construct and were incubated for 48 hr. After the incubation, the cell culture conditioned media (supernatant) from the T-150 flasks was collected and filtered through 0.45 µm pore size filter (9914-2504, Cytiva) for cultured neuron infection. The remaining HEK293T cells in the flasks were lifted using a cell scraper and were pelleted at 800 g for 10 min for making the purified and concentrated virus. Pelleted cells were then resuspended using 20 mL of 100 mM TBS (100 mM NaCl, 20 mM Tris, pH 8.0). The surfactant, 10% sodium deoxycholate (D5670-25G, Sigma-Aldrich), was then added to the resuspended pellet to a final concentration 0.5%, along with benzonase nuclease (E1014-5KU, Sigma-Aldrich) to a final concentration of 50 units/mL and incubated for 1 hour at 37°C. Cell debris were removed by centrifugation at 2500 g for 15 minutes and the supernatant was harvested. The clarified cell lysate was loaded to a pre-equilibrated HiTrap heparin column (GE17-0406-01, Cytiva) followed by a 10 mL 100 mM TBS wash using a peristaltic pump, and then successive washes of 1 mL of 200 mM TBS (200 mM NaCl, 20 mM Tris, pH 8.0), and 1 mL of 300 mM TBS (300 mM NaCl, 20 mM Tris, pH 8.0), using a syringe. The bound virus in the column was eluted using a sequence of 1.5 mL 400 mM TBS, 3 mL 450 mM TBS, and 1.5 mL 500 mM TBS (all with 20mM Tris, pH 8.0). Eluted viruses were then concentrated by centrifugation at 2000 g for 2 minutes using a 15 mL centrifugal unit with 100K molecular weight cut off (UFC910024, Sigma-Aldrich) to a final volume of 500 µL. Further concentration of viruses was achieved using a 0.5 mL centrifugal unit (UFC510024, Sigma-Aldrich) to a final volume of 100-200 µL. Titration of the concentrated viruses was performed by AAVpro Titration Kit (for Real Time PCR) Ver.2 (6233, Takara Bio Inc) following the manufacture’s protocol. The standard curve was prepared using the positive control solution provided by Takara with serial dilution for absolute quantification. The titer of each viral sample was calculated in reference to the standard curve.

### Primary neuronal culture and infection

One day prior to neuron dissociation, 24 well plates and 35 mm glass bottom dishes (D35-14-1- N, Cellvis) were coated with 0.1 mg/mL Poly-D-Lysine (PDL; P6407-5MG, Millipore) dissolved in 1X Borate buffer (28341, Thermo Scientific) overnight at room temperature. Plates and dishes were washed four times with sterile diH2O and dried in the tissue culture hood on the day of neuron dissociation. E18 rat hippocampal tissue (SDEHP, Brain Bits) were dissociated with papain (PAP, Brain Bits) and DNase I (07469, Stemcell Technologies) following the manufacturer instructions for neuron dissociation. Dissociated neurons were plated in PDL-coated plates and dishes and cultured in complete Neurobasal Plus medium (A3582901, Life Technologies) supplemented with 0.01% (v/v) Gentamicin (15710064, Life Technologies), 0.75% (v/v) Glutamax (35050061, Life Technologies), 2% (v/v) B27 Plus (A3582801, Life Technologie), 5% (v/v) FBS (F1051-500ML, Sigma-Aldrich) at 37°C, 5% CO2. The entire culture medium of each well and dish was removed 24 hr after the plating and replaced with complete Neurobasal Plus medium supplemented with 0.01% (v/v) Gentamicin, 0.75% (v/v) Glutamax, 2% (v/v) B27 Plus to stop glial growth. Subsequently, ∼50% of the media in each well was replaced every 3-4 days. At DIV6, neurons were infected with a mixture of crude supernatant of CaST-encoded AAV1/2 viruses by replacing half of the culture media. The plate and dishes were incubated for another ∼two weeks in the incubator prior to stimulation.

### Primary neuron CaST experiments

Rat hippocampal neurons were dissociated, plated, and infected as described above. At DIV17- 22 (∼two weeks after infection), neurons were ready for stimulation. Neuron stimulation reagents including Potassium Chloride (KCl; P3911-25G, Sigma-Aldrich), 2,5-dimethoxy-4- iodoamphetamine (DOI; D101, Sigma-Aldrich), and dopamine (DA; H8502, Sigma-Aldrich) were used to introduce neuron firing. A portion of the culture media from each well was taken out and saved in microcentrifuge tubes. The stimulation mixtures were generated by adding the neuron stimulation reagent and biotin solution to the saved culture media to the desired concentration. The stimulation mixtures were then added back to each well and the neurons were incubated at 37°C for 30 min for CaST labeling. For +KCl conditions, we used a final concentration of 30 mM KCl. For +DOI conditions, we used 10 µM of DOI as the final concentration. For +DA conditions, we used a final concentration of 10 µM DA. For +biotin conditions, we used a final concentration of 50 μM biotin. Neurons were then fixed and stained as described above for HEK293T cell experiments. Time series experiments were performed by treating the neurons with or without 30 mM KCl for 10 min or 30 min at 37°C. Neurons were then fixed and stained as described above for HEK293T cell experiments prior to imaging.

### Immunofluorescence imaging

Cells and neurons were imaged immediately after fixation and staining. Fluorescence images were taken with Keyence BZ-X810 fluorescence microscope (acquisition software v1.1.2) with an 80W metal halide lamp as the fluorescent light source and a PlanApo 10X air objective lens (NA 0.45). Expression of CaST was visualized by GFP using 470/40 nm excitation filter and 525/50 nm emission filter, and Alexa Fluor568 using 545/25 nm excitation filter and 605/70 nm emission filter. Biotinylated proteins were labeled by SA-647 and visualized by 620/60 nm excitation filter and 700/75 nm emission filter. Images were analyzed by custom scripts in Fiji/ImageJ v2.9.0 and MATLAB vR2020b (see “***Analysis of CaST immunofluorescence***”).

Confocal imaging was performed using a Carl Zeiss LSM 800 confocal microscope equipped with 488, 561, and 640nm lasers, and a 63x oil immersion objective (NA 1.4). 35 mm glass bottom dishes with a 14 mm micro-well and #1.5 cover glass (0.16mm-0.19mm) were used for high resolution confocal imaging. Fluorescence images were collected with a 512 x 512-pixel resolution and with a pixel dwell time of 1.03 μsec/pixel. Images were acquired using a PMT detector and emission filter ranges of 450–575 nm, 450–640 nm, and 645–700 nm for EGFP, Alexa Fluor568, and Alexa Fluor647 detection, respectively, for best signal. All images were collected and processed using ZEN software v2.3 (Carl Zeiss).

### Analysis of CaST immunofluorescence

For CaST fluorescence characterization, we analyzed all cells in the FOV that expressed CaST (assessed using the GFP channel). To accomplish this, the GFP images across all conditions of a given experiment (calcium-treated and non-treated) were pseudo combined into one super FOV, so that the exact same cell detection threshold was applied equally to all images (using the *cell-segm* automated thresholding script^61^). This ensured no bias in the detection of GFP+ cells across conditions. The cell masks were then applied to the original GFP and SA-647 images, so that the GFP and SA-647 cell fluorescence could be calculated for cells belonging to an individual FOV’s.

To ensure reproducibility, typically 6-12 FOVs were imaged for a given experimental condition. We reported both the raw GFP and raw SA-647 fluorescence of all cells pooled across all FOVs in a scatter plot. These raw scatter plots illustrate that the SA-647 labeling in calcium-treated cells is higher than the SA-647 labeling observed in untreated cells across generally all GFP expression levels of the tool (the entire x-axis).

To make a quantitative comparison of this data that matched the standards previously reported for evaluating similar tools (such as FLARE^5^, FLiCRE^6^, and Cal-Light^7^), we calculated the mean SA-647/GFP cell ratios found for each FOV. This analysis can normalize for any difference in expression levels of the tool across cells and conditions. Matching the standards of these previously published works, the background SA-647 autofluorescence in the epifluorescence images was subtracted from every cell per FOV prior to taking the ratio to the GFP fluorescence values (autofluorescence was calculated as the mean pixel value of the entire image, excluding pixels that corresponded to cell masks). See **Supplementary** Figure 2 for an example of each step of this analysis pipeline, along with appropriate summary data.

### Neuron viability analysis

Neuron viability assays were performed using DRAQ7 dye (D15105, Invitrogen). Rat hippocampal neurons were dissociated, plated as described above and were infected with crude supernatant AAV1/2 viruses for either the complete CaST, or only one CaST fragment (CD4- sTb(C)-M13-GFP) as a DRAQ7- negative control, at DIV6. At DIV19, neurons were stained with DRAQ7 solution with a final concentration of 3 µM for 10 mins at 37°C, protected from light. After staining, neurons were washed with DPBS, fixed with 4% PFA, and imaged in DPBS. For DRAQ7+ control, neurons expressing complete CaST constructs were fixed with 4% PFA and were permeabilized with methanol for 8 mins at -20°C at DIV19. Neurons were then incubated with DRAQ7 dye solution for 10 mins at 37°C, protected from light and were washed and imaged using fluorescence microscopy.

### Calcium imaging in cultured neurons

Rat hippocampal neurons were dissociated, plated as described above and were infected with a mixture of crude supernatant of CaST and RCaMP2 encoded AAV1/2 viruses by replacing half of the culture media at DIV6. The plate and dishes were incubated for another ∼two weeks in the incubator prior to stimulation. For simultaneous RCaMP2 recording and CaST labeling characterization, mild neuronal stimulation was introduced by replacing half volume of the neuron culture media and biotin was also introduced at a final concentration of 50 µM for CaST labeling. Calcium activity of each RCaMP2 positive neuron was continuously recorded for 5 min post-stimulation using the Keyence microscope. Neurons were treated with biotin for a total of 30 min at room temperature. After stimulation, neurons were washed, fixed, and stained as described above. The same FOV recorded during RCaMP2 imaging was then re-identified, and images showing CaST expression and labeling were captured.

For calcium imaging during drug-treatment, neurons were infected with RCaMP2 encoded AAV1/2 virus at DIV6. The neurons were incubated for another ∼two weeks in the incubator prior to stimulation. KCl, DA, and DOI stimulations were induced as described above. Calcium activity of each RCaMP2 positive neuron was continuously recorded for 1 min pre- and 1 min post-stimulation using the Keyence microscope.

The mean RCaMP2 FOV during the recording was input to Cellpose^62^ (v2.2.3) to identify masks corresponding to individual neurons. Pixels corresponding to individual cell masks were then analyzed in Fiji to obtain the fluorescence time series of all neurons in the FOV, and values were imported into MATLAB vR2020b for further analysis. The time series of each neuron was reported either as dF/F (**Extended Data** Figure 6), or reported as a Z-score relative to the baseline recording (**Extended Data** Figure 8). dF/F was calculated as (Ft-Fb)/Fb, where Fb = 2^nd^ percentile of each cell’s fluorescence timeseries. Z-scores were calculated in MATLAB using the “zscore” function. To quantify the mean RCaMP2 activity following mild stimulation, the “findpeaks” function was used in MATLAB, and the detected peak heights was averaged for each neuron’s activity trace (**Extended Data** Figure 6). The quantify the mean peak height following the drug-induced baseline increase following calcium influx, the maximum of each cell’s calcium trace during the post-stimulation period was reported (**Extended Data** Figure 8).

### Optogenetic stimulation for CaST labeling in cultured neurons

Rat hippocampal neurons were dissociated, plated as described above and were infected with a mixture of crude supernatant of CaST and bReaChES encoded AAV1/2 viruses by replacing half of the culture media at DIV6 and were immediately wrapped in foil. At DIV18, dishes with CaST and bReaChES co-expressing rat hippocampal neurons were covered on the bottom with black tape to block the orange stimulation light (M595F2, Thorlabs, 5.67 mw/mm^2^). A ∼1-mm slit was left open allowing spatially targeted bReaChES stimulation. Biotin was introduced at a final concentration of 50 µM for CaST labeling. Orange light was cycled every 6.5 seconds with 2 seconds on and 4.5 seconds off. The total stimulation was 30 minutes long, using 5-ms long pulses delivered at 20Hz during the “on” cycle. Glutamate receptor antagonist APV and NBQX were added at a final concentration of 50 µM and 20 µM respectively at the time of light stimulation to reduce synchronized neuron firing across the entire dish. After stimulation, neurons were washed, fixed, and stained as described above. Images were taken using Keyence BZ-X810 with tile scan mode.

### Mouse animal models

All experimental and surgical protocols were approved by the University of California, Davis, Institutional Animal Care and Use Committee. For CaST experiments, 5-7 week old male and female wild-type C57BL/6J (Jackson Laboratory Strain 000664) mice were used. Mice were maintained on a 12h reverse light-dark cycle (lights on at 9PM) at 22⁰C and 40-60% humidity, group-housed with same-sex cage mates and given ad-lithium access to food and water.

### Mouse stereotaxic surgeries

Briefly, mice were maintained under anesthesia with 1.5 – 2% isoflurane and placed in a stereotaxic apparatus (RWD) on a heating pad. The fur on the top of the skull was removed and antiseptic iodine and 70% alcohol were used in alternation to clean the scalp. Sterile ocular lubricant (Dechra) was administered to the eyes of the mice to protect them from drying out. A midline scalp incision was made, and 0.1% hydrogen peroxide was applied to the skull. A craniotomy was made above the injection site. Virus was then injected into the targeted region using a 33 gauge beveled needle (WPI) and a 10 μL Hamilton syringe controlled by an injection pump (WPI). For all surgeries, 1000 nL total of virus were injected into the targeted mPFC brain region (coordinates: ML +/- 0.5, AP +1.98, DV -2.25) at a rate of 150 nL min^-1^.

For 2-photon imaging surgeries, 1000 nL of AAV5-CaMKIIa-GCaMP6f was injected into mPFC (diluted 1:1 in DPBS; Addgene, 100834-AAV5, 2.2e12 VG/mL titer). A 1-mm diameter GRIN lens (Inscopix) was then implanted above the mPFC (ML +/- 0.5, AP +1.98, DV -2.05). Implants and custom stainless steel headplates were secured to the skull using dental adhesive and cement system (Pentron, C&B metabond). For CaST surgeries, 500 nL of a 1:1 ratio of the two home-made CaST viruses was injected into mPFC (AAV2/1-Syn-CD4-sTb(C)-M13-GFP, 4.85e9 copies/mL; AAV2/1-Syn-CaM-V5-sTb(N), 4.79e9 copies/mL). Once the surgery was completed, the incision was closed with tissue adhesive (GLUture). Mice received a dose of 3.25 mg/kg EthiqaXR for pain recovery and was revived in a new, clean cage placed on a heating pad.

### 2-photon Ca^2+^ imaging in mice

2-photon Ca^2+^ imaging was performed using a commercial microscope (2P+, Bruker) and a 16x, 0.8 NA objective (MRP07220, Nikon). A tunable IR femtosecond pulse laser set to 920 nm (Coherent, Discovery TPC) was used for excitation, and fluorescence emission was collected using a GaAsP PMT (H10770PB-40, Hamamatsu). The excitation laser was directed by galvo scanners sampling 512x512 pixels. Each image was captured at 2 Hz. The imaging field of view was 448x448 μm (optical zoom of 2.5x). Data were collected using the PrairieView v5.6 software and analyzed using Suite2P^63^ (v0.14.3) and custom MATLAB vR2020b scripts. Fluorescence values corresponding to each cell mask output from Suite2P were then averaged to create a fluorescence time series for each neuron. Each neuron’s trace was z-scored (“zscore” function in MATLAB) and smoothed using a 5-s sliding window (“smooth” function in MATLAB). We calculated the difference between each neuron’s 10-minute psilocybin activity trace and 10-minute saline activity trace (“Psilocybin minus Saline” activity trace). We then took the average of this difference and ranked cells according to this “Avg diff” value. We considered “Activated” neurons to be those with an Avg diff Z-score value > 0.05, and “inhibited” neurons to be those with an Avg diff Z-score value < -0.05.

### CaST experiments in mice

Mice were handled and given USP grade saline (.9%) injections for three consecutive days before biotin labeling. After a week of viral expression, experimental mice were given two injections, one 24 mg/kg intraperitoneal (IP) injection of diluted biotin solution (B4639, Sigma-Aldrich; dissolved in DMSO as a 10 mM stock solution) and one 3 mg/kg IP injection of psilocybin (synthesized as previously described^36^) dissolved in saline. Control mice were injected with biotin solution and saline (5 ml/kg IP). Mice were placed in separated clean cages following the injection and were sacrificed 1 hour after IP injections. A subset of mice was video recorded for head twitch data acquisition following injections. The number of HTRs during the first 20 minutes of video recording was quantified, manually scored by a blinded experimenter.

After biotin labeling, mice were perfused with ice-cold PBS, followed by 4% PFA, and brains were collected and stored in 4% PFA overnight at 4°C. The next day, brains were switched out from PFA and stored in PBS until slicing. 60 μm slices were collected from the mPFC and placed in wells with PBS and stored in 4°C. Slices were washed with PBS-T for 2 mins (3x), then blocked in 5% Normal Donkey Serum (017-000-121, Jackson ImmunoResearch) and 0.3% Triton-X (T9284-100ML, Sigma-Aldrich; in PBS-T) for 1 hour at room temperature. Slices were stained with streptavidin-Alexa647 (1:1000 dilution; S32357, Invitrogen) in 5% NDS/PBS-T for 1.5 hours at room temperature. Slices were washed with PBS-T for 5 mins at room temperature and then mounted with DAPI-Fluoromount-G (0100-20, SouthernBiotech) to adhere coverslips. 40x images were taken on BZ-810 Keyence fluorescence microscope and were analyzed as described above for “***Analysis of CaST immunofluorescence”*.**

### c-Fos staining of mouse brain slices

Mice were injected with CaST and treated with saline or psilocybin as described above. Following perfusion of the brain and slicing, a subset of slices were saved for c-Fos only staining. Slices were washed in PBST (1% Triton in PBS) for 2 hours, placed on shaker at room temperature. Slices were blocked in 1% BSA/PBST for 1 hour at room temperature, and then stained with 300 µL of rat anti-cFos primary antibody (226017, Synaptic Systems) at 1:1000 dilution in 1% BSA/PBST overnight on a shaker at room temperature. The next day, slices were washed with PBST for 20 mins, three times, placed on a shaker at room temperature. Slices were then stained with 300 µL of anti**-**rat-Alexa568 secondary antibody (A78946, Invitrogen) at 1:500 dilution in 1% BSA/PBST for 2 hours on a shaker at room temperature. Slices were washed with PBST for 20 min, two times, placed on a shaker at room temperature. Mounted slices and used DAPI-Fluoromount-G to adhere coverslips. For simultaneous CaST staining, the same procedure was followed, except streptavidin-Alexa647 was also included during the secondary antibody incubation (1:1000 dilution; S32357, Invitrogen). 40x images were taken on BZ-810 Keyence fluorescence microscope, and were analyzed using an automated cell-detection method in MATLAB (cellsegm)^61^ for each channel individually.

### Summary of statistical analyses

Statistical analysis including Mann-Whitney U test, ordinary one-way ANOVA, two-way ANOVA, Pearson correlation analysis, and Wilson/Brown ROC curve analysis were performed in GraphPad Prism v9.0 (GraphPad Software). The D’Agostino & Pearson test for normality was performed in Prism prior to using any parametric statistical tests. Significance was defined as a \**P* < 0.05, \*\**P* < 0.01, and \*\*\**P* < 0.001 for the defined statistical test (n.s., not significant *P* ≥ 0.05). All experiments performed in study were independently replicated at least twice.

## DATA AVAILABILITY

Plasmids and associated DNA sequences generated in this study are available on Addgene (catalog nos. 219779–219784; https://www.addgene.org/Christina_Kim/). There are no restrictions on data availability. Source data are provided with this paper.

## CODE AVAILABILITY

Custom MATLAB scripts used to analyze images are freely available at https://github.com/tinakimlab/CaST-analysis-scripts.

## REFERENCES

1. Grynkiewicz, G., Poenie, M. & Tsien, R.Y. A new generation of Ca2+ indicators with greatly improved fluorescence properties. Journal of biological chemistry 260, 3440–3450 (1985).

2. Tian, L. et al. Imaging neural activity in worms, flies and mice with improved GCaMP calcium indicators. Nature methods 6, 875–881 (2009).

3. Chen, T.-W. et al. Ultrasensitive fluorescent proteins for imaging neuronal activity. Nature 499, 295–300 (2013).

4. Inoue, M. et al. Rational design of a high-affinity, fast, red calcium indicator R-CaMP2. Nature methods 12, 64–70 (2015).

5. Wang, W. et al. A light- and calcium-gated transcription factor for imaging and manipulating activated neurons. Nature Biotechnology 35, 864–871 (2017).

6. Kim, C.K. et al. A Molecular calcium integrator reveals a striatal cell type driving aversion. Cell 183, 2003–2019. e2016 (2020).

7. Lee, D., Hyun, J.H., Jung, K., Hannan, P. & Kwon, H.-B. A calcium- and light-gated switch to induce gene expression in activated neurons. Nature Biotechnology 35, 858–863 (2017).

8. Fosque, B.F. et al. Labeling of active neural circuits in vivo with designed calcium integrators. Science 347, 755–760 (2015).

9. Motta-Mena, L.B. et al. An optogenetic gene expression system with rapid activation and deactivation kinetics. Nature chemical biology 10, 196–202 (2014).

10. McKinney, S.A., Murphy, C.S., Hazelwood, K.L., Davidson, M.W. & Looger, L.L. A bright and photostable photoconvertible fluorescent protein. Nature Methods 6, 131–133 (2009).

11. Allen, W.E. et al. Thirst-associated preoptic neurons encode an aversive motivational drive. Science 357, 1149–1155 (2017).

12. Reijmers, L.G., Perkins, B.L., Matsuo, N. & Mayford, M. Localization of a Stable Neural Correlate of Associative Memory. Science 317, 1230–1233 (2007).

13. Liu, X. et al. Optogenetic stimulation of a hippocampal engram activates fear memory recall. Nature 484, 381–385 (2012).

14. Choi, J.-E., Kim, J. & Kim, J. Capturing activated neurons and synapses. Neuroscience Research 152, 25–34 (2020).

15. Gore, F. et al. Neural representations of unconditioned stimuli in basolateral amygdala mediate innate and learned responses. Cell 162, 134–145 (2015).

16. Cho, K.F. et al. Split-TurboID enables contact-dependent proximity labeling in cells. Proceedings of the National Academy of Sciences 117, 12143–12154 (2020).

17. Roux, K.J., Kim, D.I., Raida, M. & Burke, B. A promiscuous biotin ligase fusion protein identifies proximal and interacting proteins in mammalian cells. Journal of Cell Biology 196, 801–810 (2012).

18. Branon, T.C. et al. Efficient proximity labeling in living cells and organisms with TurboID. Nature biotechnology 36, 880–887 (2018).

19. Spector, R. & Mock, D. Biotin transport through the blood-brain barrier. Journal of neurochemistry 48, 400–404 (1987).

20. Bongarzone, S. et al. Imaging biotin trafficking in vivo with positron emission tomography. Journal of medicinal chemistry 63, 8265–8275 (2020).

21. Kim, J.H. et al. High cleavage efficiency of a 2A peptide derived from porcine teschovirus-1 in human cell lines, zebrafish and mice. PloS one 6, e18556 (2011).

22. Mountford, P.S. & Smith, A.G. Internal ribosome entry sites and dicistronic RNAs in mammalian transgenesis. Trends in Genetics 11, 179–184 (1995).

23. Licursi, M., Christian, S., Pongnopparat, T. & Hirasawa, K. In vitro and in vivo comparison of viral and cellular internal ribosome entry sites for bicistronic vector expression. Gene therapy 18, 631–636 (2011).

24. Kim, C.K., Cho, K.F., Kim, M.W. & Ting, A.Y. Luciferase-LOV BRET enables versatile and specific transcriptional readout of cellular protein-protein interactions. eLife 8, e43826 (2019).

25. Kim, C.K. et al. Simultaneous fast measurement of circuit dynamics at multiple sites across the mammalian brain. Nature methods 13, 325–328 (2016).

26. Gasbarri, A., Sulli, A. & Packard, M.G. The dopaminergic mesencephalic projections to the hippocampal formation in the rat. Progress in Neuro-Psychopharmacology and Biological Psychiatry 21, 1–22 (1997).

27. Yu, Q. et al. Genetic labeling reveals temporal and spatial expression pattern of D2 dopamine receptor in rat forebrain. Brain Structure and Function 224, 1035–1049 (2019).

28. Bombardi, C. Neuronal localization of 5-HT2A receptor immunoreactivity in the rat hippocampal region. Brain research bulletin 87, 259–273 (2012).

29. Gurevich, E.V., Gainetdinov, R.R. & Gurevich, V.V. G protein-coupled receptor kinases as regulators of dopamine receptor functions. Pharmacological research 111, 1–16 (2016).

30. McCorvy, J.D. & Roth, B.L. Structure and function of serotonin G protein-coupled receptors. Pharmacology & therapeutics 150, 129–142 (2015).

31. Neve, K.A., Seamans, J.K. & Trantham-Davidson, H. Dopamine receptor signaling. Journal of receptors and signal transduction 24, 165–205 (2004).

32. Dhyani, V. et al. GPCR mediated control of calcium dynamics: A systems perspective. Cellular signalling 74, 109717 (2020).

33. Ly, C. et al. Psychedelics promote structural and functional neural plasticity. Cell reports 23, 3170–3182 (2018).

34. Vargas, M.V. et al. Psychedelics promote neuroplasticity through the activation of intracellular 5-HT2A receptors. Science 379, 700–706 (2023).

35. Shao, L.-X. et al. Psilocybin induces rapid and persistent growth of dendritic spines in frontal cortex in vivo. Neuron 109, 2535–2544. e2534 (2021).

36. Cameron, L.P. et al. 5-HT2ARs Mediate Therapeutic Behavioral Effects of Psychedelic Tryptamines. ACS Chemical Neuroscience (2023).

37. Catlow, B.J., Song, S., Paredes, D.A., Kirstein, C.L. & Sanchez-Ramos, J. Effects of psilocybin on hippocampal neurogenesis and extinction of trace fear conditioning. Experimental brain research 228, 481–491 (2013).

38. Hibicke, M., Landry, A.N., Kramer, H.M., Talman, Z.K. & Nichols, C.D. Psychedelics, but not ketamine, produce persistent antidepressant-like effects in a rodent experimental system for the study of depression. ACS chemical neuroscience 11, 864–871 (2020).

39. Hesselgrave, N., Troppoli, T.A., Wulff, A.B., Cole, A.B. & Thompson, S.M. Harnessing psilocybin: antidepressant-like behavioral and synaptic actions of psilocybin are independent of 5-HT2R activation in mice. Proceedings of the National Academy of Sciences 118, e2022489118 (2021).

40. Cameron, L.P. et al. A non-hallucinogenic psychedelic analogue with therapeutic potential. Nature 589, 474–479 (2021).

41. Corne, S., Pickering, R. & Warner, B. A method for assessing the effects of drugs on the central actions of 5-hydroxytryptamine. British journal of pharmacology and chemotherapy 20, 106–120 (1963).

42. Halberstadt, A.L. & Geyer, M.A. Characterization of the head-twitch response induced by hallucinogens in mice: detection of the behavior based on the dynamics of head movement. Psychopharmacology 227, 727–739 (2013).

43. González-Maeso, J. et al. Hallucinogens recruit specific cortical 5-HT2A receptor- mediated signaling pathways to affect behavior. Neuron 53, 439–452 (2007).

44. Jefsen, O.H., Elfving, B., Wegener, G. & Müller, H.K. Transcriptional regulation in the rat prefrontal cortex and hippocampus after a single administration of psilocybin. Journal of Psychopharmacology 35, 483–493 (2021).

45. Davoudian, P.A., Shao, L.-X. & Kwan, A.C. Shared and distinct brain regions targeted for immediate early gene expression by ketamine and psilocybin. ACS Chemical Neuroscience 14, 468–480 (2023).

46. Wood, J., Kim, Y. & Moghaddam, B. Disruption of prefrontal cortex large scale neuronal activity by different classes of psychotomimetic drugs. Journal of Neuroscience 32, 3022–3031 (2012).

47. Michaiel, A.M., Parker, P.R. & Niell, C.M. A hallucinogenic serotonin-2A receptor agonist reduces visual response gain and alters temporal dynamics in mouse V1. Cell reports 26, 3475–3483. e3474 (2019).

48. Ziv, Y. et al. Long-term dynamics of CA1 hippocampal place codes. Nature neuroscience 16, 264 (2013).

49. Dong, C. et al. Psychedelic-inspired drug discovery using an engineered biosensor. Cell 184, 2779–2792. e2718 (2021).

50. Rayaprolu, S. et al. Cell type-specific biotin labeling in vivo resolves regional neuronal and astrocyte proteomic differences in mouse brain. Nature Communications 13, 2927 (2022).

51. Chan, K.Y. et al. Engineered AAVs for efficient noninvasive gene delivery to the central and peripheral nervous systems. Nature neuroscience 20, 1172–1179 (2017).

52. Chen, K.H., Boettiger, A.N., Moffitt, J.R., Wang, S. & Zhuang, X. Spatially resolved, highly multiplexed RNA profiling in single cells. Science 348, aaa6090 (2015).

53. Wang, X. et al. Three-dimensional intact-tissue sequencing of single-cell transcriptional states. *Science*, eaat5691 (2018).

54. Lim, M.J. et al. MALDI HiPLEX-IHC: multiomic and multimodal imaging of targeted intact proteins in tissues. Frontiers in Chemistry 11, 1182404 (2023).

55. Stoeckius, M. et al. Simultaneous epitope and transcriptome measurement in single cells. Nature methods 14, 865–868 (2017).

56. Mirdita, M. et al. ColabFold: making protein folding accessible to all. Nature methods 19, 679–682 (2022).

57. Jumper, J. et al. Highly accurate protein structure prediction with AlphaFold. Nature 596, 583–589 (2021).

58. Feinberg, E.H. et al. GFP Reconstitution Across Synaptic Partners (GRASP) defines cell contacts and synapses in living nervous systems. Neuron 57, 353–363 (2008).

59. Hanke, T., Szawlowski, P. & Randall, R.E. Construction of solid matrix-antibody-antigen complexes containing simian immunodeficiency virus p27 using tag-specific monoclonal antibody and tag-linked antigen. Journal of general virology 73, 653–660 (1992).

## METHODS-ONLY REFERENCES

60. Konermann, S. et al. Optical control of mammalian endogenous transcription and epigenetic states. Nature 500, 472–476 (2013).

61. Hodneland, E., Kögel, T., Frei, D.M., Gerdes, H.-H. & Lundervold, A. CellSegm-a MATLAB toolbox for high-throughput 3D cell segmentation. Source code for biology and medicine 8, 16 (2013).

62. Stringer, C., Wang, T., Michaelos, M. & Pachitariu, M. Cellpose: a generalist algorithm for cellular segmentation. Nature methods 18, 100–106 (2021).

63. Pachitariu, M. et al. Suite2p: beyond 10,000 neurons with standard two-photon microscopy. BioRxiv, 061507 (2016).

