## Extended Data Figs 1-9 for "Rapid, biochemical tagging of cellular activity history in vivo"

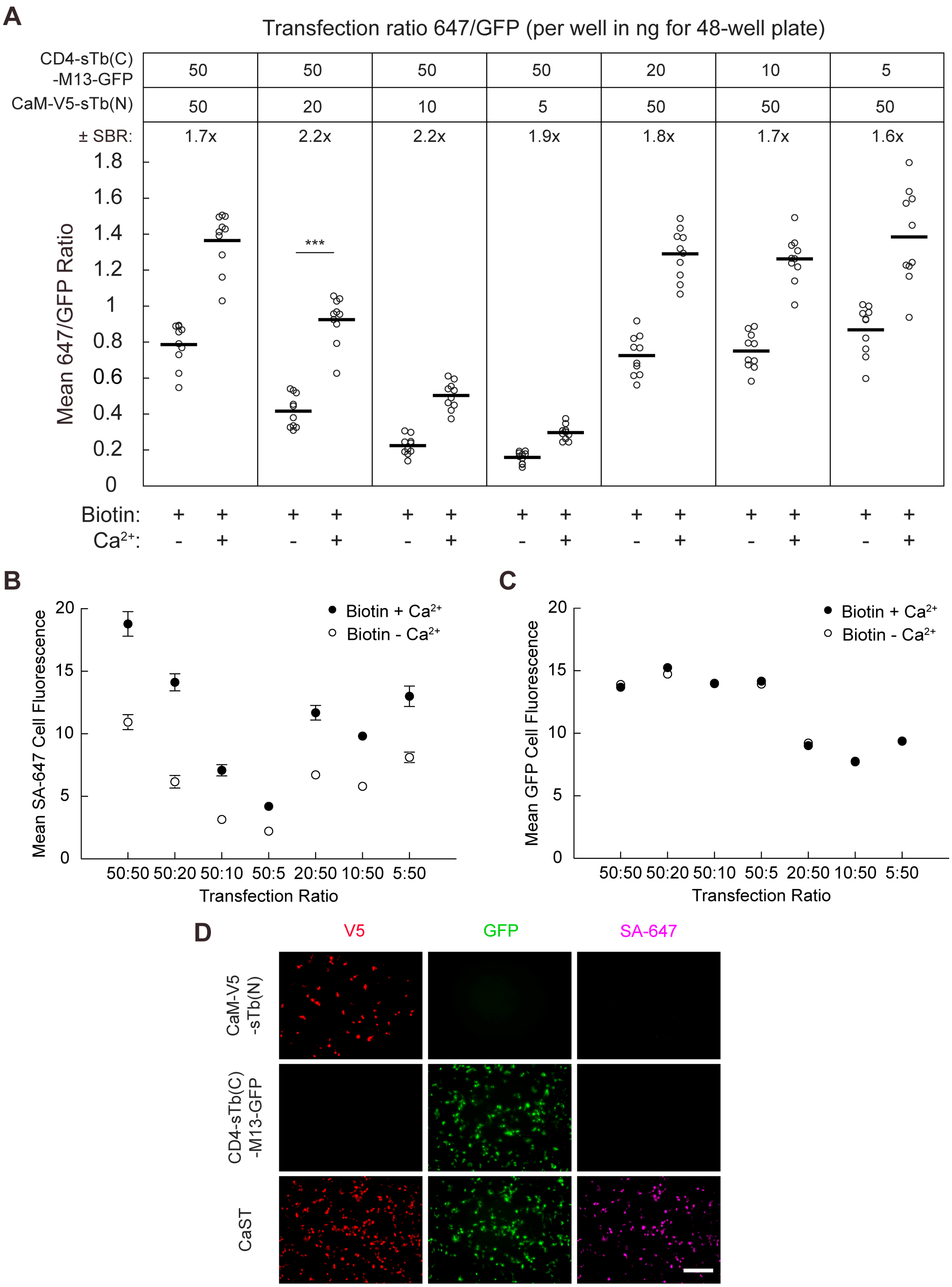


**Extended Data Figure 1. Comparison of different CaST transfection ratios of fragments.**
**A)** HEK cells were transfected with different ratios of the CaST fragments indicated above in ng for 48-well plate and incubated overnight. Cells were treated with 50 μM biotin and ± $Ca^{2+}$ (5

mM $CaCl_2$ and 1 µM ionomycin) for 30 minutes. The ± $Ca^{2+}$ SBR and the FOV averages of the SA647/GFP fluorescence ratio per cell are shown. The ratio of 50:20 (CD4-sTb(C)-M13-GFP:CaM-V5-sTb(N)) had a significant 2.2-fold ± $Ca^{2+}$ SBR ($n$ = 10 FOVs per condition; $P$ = 1.1e-5, U = 0, two-tailed Mann-Whitney U test).
**B,C)** The FOV average of the SA-647 cell fluorescence (**B**) and the GFP cell fluorescence (**C**) was calculated for each transfection ratio (CD4-sTb(C)-M13-GFP:CaM-V5-sTb(N)) after biotin ± $Ca^{2+}$ stimulation ($n$ = 10 FOVs per condition). Data are plotted as mean ± s.e.m.
**D)** HEK cells were transfected with either the Cam-V5-sTb(N) fragment (top), the CD4-sTb(C)-M13-GFP fragment (middle), or both fragments of CaST together (bottom). Cells were treated with 50 µM biotin and $Ca^{2+}$ (5 mM $CaCl_2$ and 1 µM ionomycin) for 30 minutes, then washed, stained, and imaged. Data were replicated across 12 FOVs. Scale bar, 300 µm. ****$P$<0.0001.

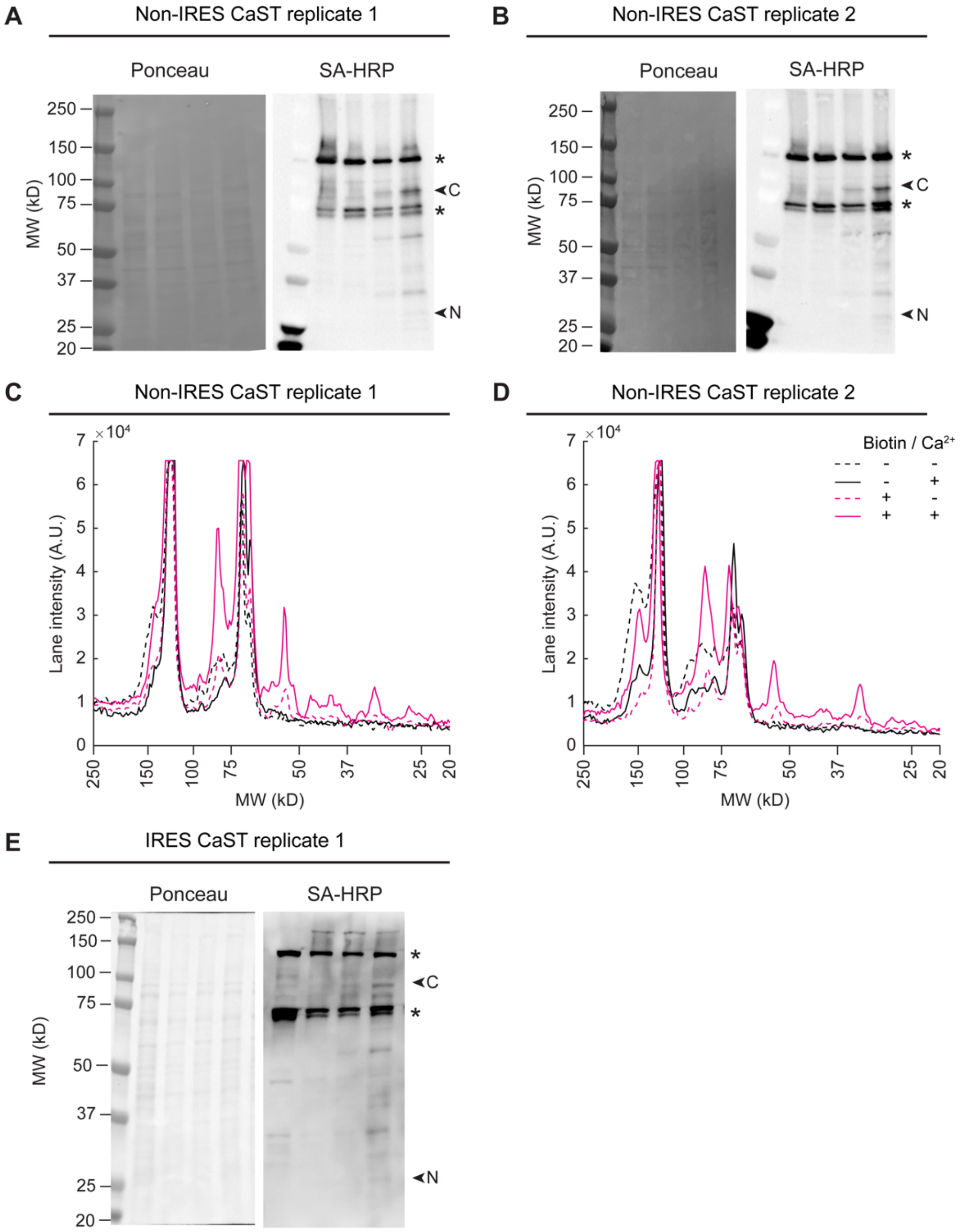


**Extended Data Figure 2. Additional western blot analysis of CaST labeling.**
**A)** Ponceau staining and raw image of the full SA-HRP blot from Figure **1C**.
**B)** Biological replicate of experiment in panel **A**.
**C,D)** Line plot profile of the entire lane of the raw blot shown in panels **A** and **B** (drawn vertically from the 250 kB to the 20 kB ladder line and centered horizontally within each lane).
**E)** Additional biological replicate experiment of panels **A-B**, except using CaST-IRES. Cells were otherwise treated identically.
*indicate bands showing endogenously biotinylated proteins. "N" indicates the expected size of the CaM-V5-sTb(N) fragment, while "C" indicates the expected size of the CD4-sTb(C)-M13-GFP fragment.

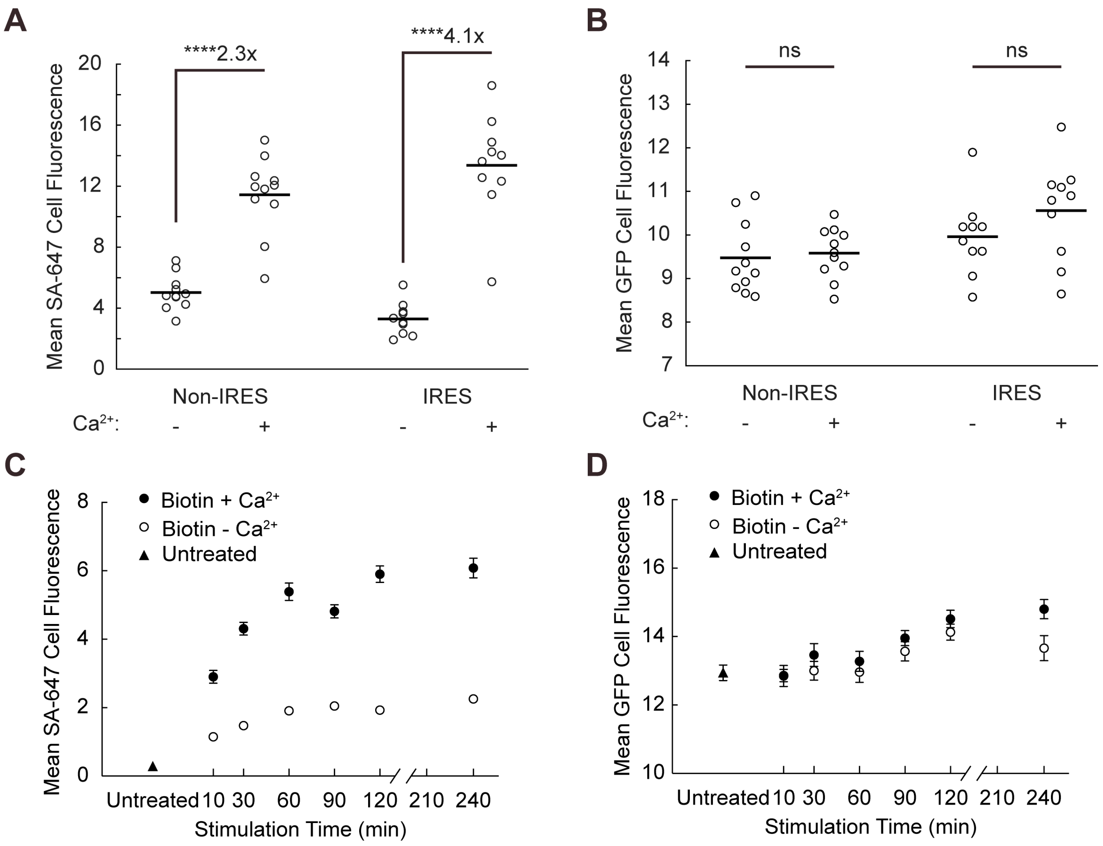


**Extended Data Figure 3. Additional characterization of CaST in HEK cells.**
**A)** The FOV averages of the SA-647 fluorescence per cell from the non-IRES data shown in **Fig. 2B**, and the IRES data shown in **Fig. 2F**. The non-IRES version exhibited a $Ca^{2+}$-dependent SBR of 2.3x (*n* = 11 FOVs per condition; *P* = 1.1e-5, U = 2, two-tailed Mann-Whitney U test). The $Ca^{2+}$-dependent SBR for the IRES version was 4.1x (*n* = 10 FOVs per condition; *P* = 1.1e-5, U = 0, two-tailed Mann-Whitney U test).
**B)** The FOV averages of the GFP cell fluorescence per cell from the non-IRES data shown in **Fig. 2B** (*n* = 11 FOVs per condition; *P* = 0.562, U = 51, two-tailed Mann-Whitney U test), and the IRES data shown in **Fig. 2F** (*n* = 10 FOVs per condition; *P* = 0.165, U = 31, two-tailed Mann-Whitney U test).
**C,D)** The FOV average of the SA-647 (**C**) or GFP (**D**) cell fluorescence was calculated for each stimulation duration for data shown in **Fig. 3E-F** (*n* = 10 FOVs per condition). Cells were transfected with CaST-IRES. The untreated condition is shown on the left. Data are plotted as mean ± s.e.m. ****$P$<0.0001, ns, not significant.

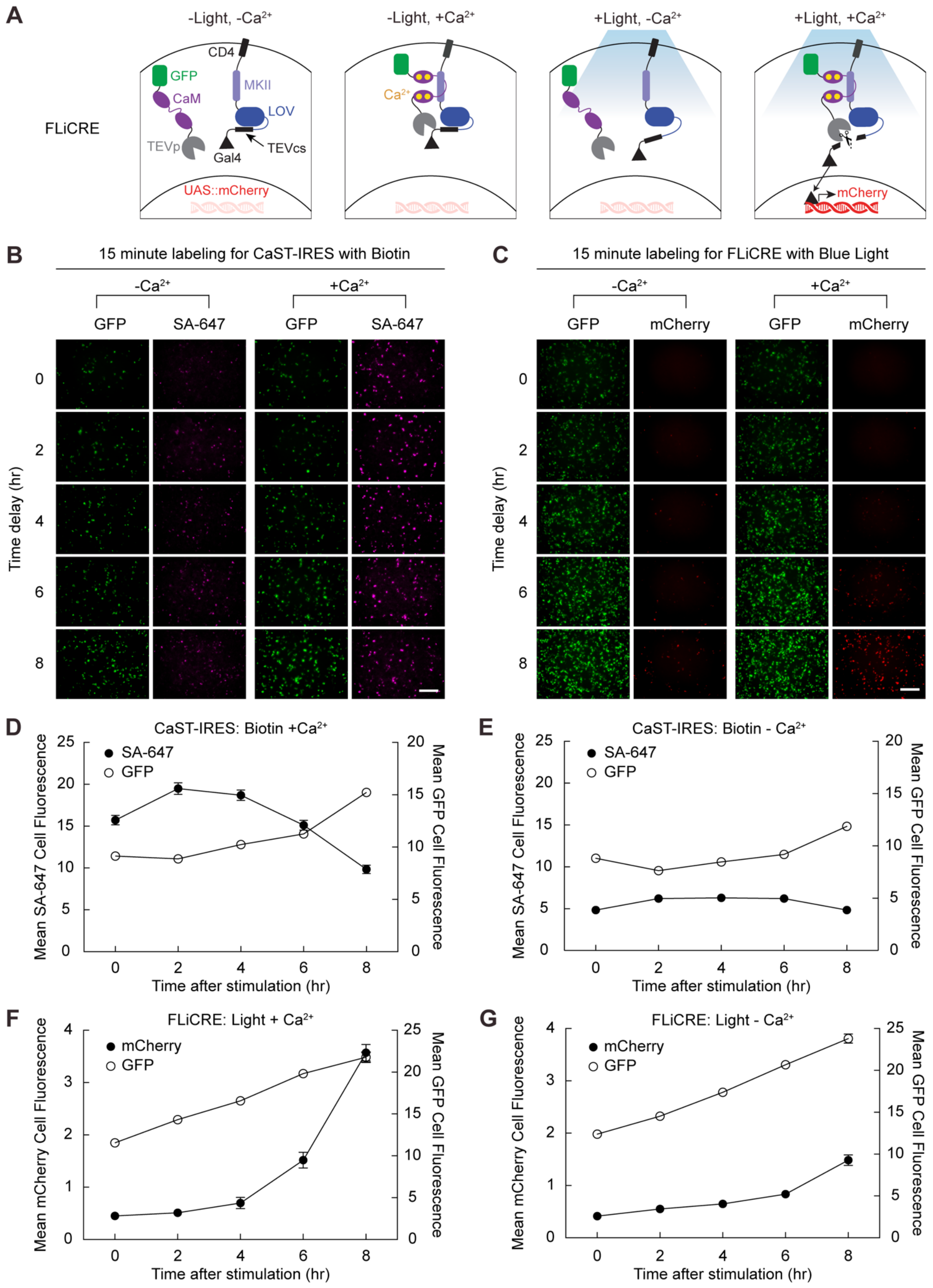

A
-Light, -Ca2+
-Light, +Ca2+
+Light, -Ca2+
+Light, +Ca2+
FLiCRE
CD4
GFP
CaM
MKII
LOV
TEVp
Gal4
TEVcs
UAS::mCherry
Ca2+
mCherry
B
15 minute labeling for CaST-IRES with Biotin
-Ca2+
+Ca2+
GFP
SA-647
Time delay (hr)
C
15 minute labeling for FLiCRE with Blue Light
GFP
mCherry
D
CaST-IRES: Biotin +Ca2+
SA-647
GFP
Mean SA-647 Cell Fluorescence
Mean GFP Cell Fluorescence
Time after stimulation (hr)
E
CaST-IRES: Biotin - Ca2+
F
FLiCRE: Light + Ca2+
mCherry
Mean mCherry Cell Fluorescence
G
FLiCRE: Light - Ca2+

**Extended Data Figure 4. Comparison of CaST to FLiCRE.**
**A)** Schematic of FLiCRE as a light- and $Ca^{2+}$-dependent transcriptional reporter. A TEV protease (TEVp) is tethered to GFP-CaM and expressed in the cytosol. A CD4-MKII-LOV-TEVcs-Gal4 fusion is expressed at the membrane. In the dark, the LOV protein cages the TEV cleavage site (TEVcs), protecting it from the TEVp. When there is high intracellular $Ca^{2+}$, CaM-M13 interact to bring the TEVp nearby the TEVcs. However, only when blue light is simultaneously delivered, will the TEVcs become uncaged and available for cleavage. With both blue light and high intracellular $Ca^{2+}$, the TEVp will cut the TEVcs, and the released Gal4 then enters the nucleus to drive expression of the UAS reporter gene.
**B,C)** Example FOVs for CaST (**B**) and FLiCRE (**C**) experiments quantified in **Fig. 4C-F**. Scale bar, 300 µm.
**D,E)** For CaST, the FOV average of the SA-647 cell fluorescence and the GFP cell fluorescence was calculated following a variable delay period after biotin + $Ca^{2+}$ (**D**) or biotin - $Ca^{2+}$ (**E**) stimulation ($n$ = 12 FOVs for conditions with 0, 4, 6, 8 hr delay time after stimulation; $n$ = 11 FOVs for conditions with 2 hr delay time after stimulation).
**F,G)** For FLiCRE, the FOV average of the UAS-mCherry cell fluorescence and the GFP cell fluorescence was calculated following a variable delay period after light + $Ca^{2+}$ (**D**) or light - $Ca^{2+}$ (**E**) stimulation ($n$ = 11 FOVs for conditions with 0 hr delay time after stimulation; $n$ = 12 FOVs for conditions with 2, 4, 6, 8 hr delay time after stimulation). Data are plotted as mean ± s.e.m. in **D**-**G**.

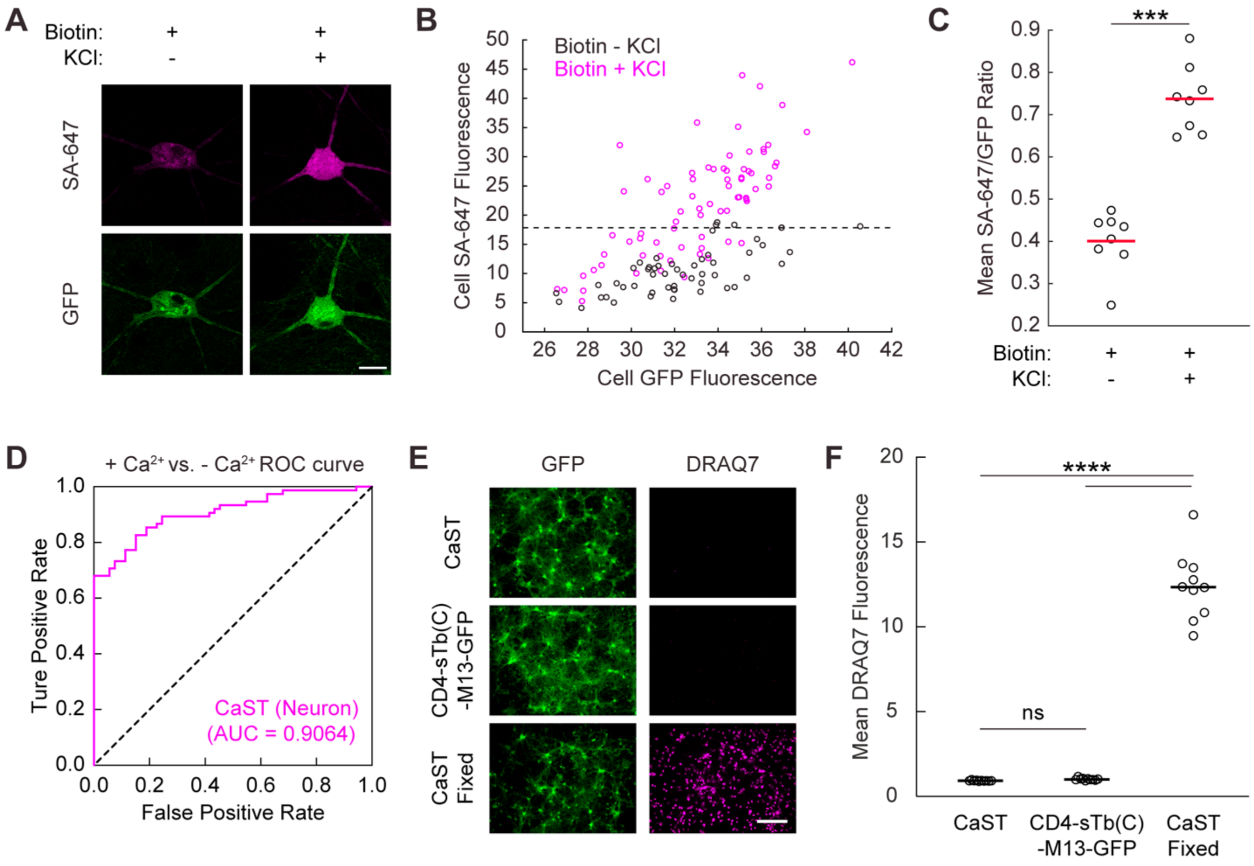


**Extended Data Figure 5. Additional characterization of CaST in neurons.**

**A)** Example confocal images of cultured rat hippocampal neurons infected with both components of CaST. Neurons were treated with 50 μM biotin ± 30 mM KCl for 30 minutes. They were then washed, fixed, and stained for SA-647. Scale bar, 20 μm.

**B)** Scatter plot of the SA-647 versus GFP fluorescence calculated for each GFP+ neuron detected across FOVs treated with biotin - KCl (*n* = 53 neurons pooled from 8 FOVs) or biotin + KCl (*n* = 75 neurons pooled from 8 FOVs). Dashed line indicates the $90^{th}$ percentile of SA-647 fluorescence values of all neurons in the biotin - KCl group.

**C)** The FOV averages of the SA-647/GFP fluorescence ratio per cell from the data shown in panel **B** (*n* = 8 FOVs per condition; *P* = 1.6e-4, U = 0, two-tailed Mann-Whitney U test).

**D)** ROC curve for distinguishing KCl-treated vs. non-treated neuron populations based on the SA-647/GFP ratios from panel **B** (AUC = 0.91, *P* = 5.5e-15, Wilson/Brown's method).

**E)** Example FOV images of CaST expressing neuron viability experiment. Neurons expressing both fragments of CaST together (top) or the CD4-sTb(C)-M13-GFP fragment (middle) were stained with DRAQ7 at a final concentration of 3 μM at DIV 19 before fixation. Neurons expressing both fragments of CaST (bottom) were fixed, permeabilized, and stained with DRAQ7 at DIV 19. Scale bar, 300 μm.

**F)** The FOV averages of the DRAQ7 fluorescence for the 3 conditions shown in panel **D** (*n* = 10 FOVs per condition; CaST versus CD4-sTb(C)-M13-GFP: *P* = 0.9831; CaST versus CaST Fixed: *P* = 8.0e-15; CD4-sTb(C)-M13-GFP versus CaST Fixed: *P* = 8.0e-15, Tukey's post-hoc multiple comparison's test following a 1-way ANOVA, $F_{2,27}$ = 326.3, *P* = 1.2e-19). ****P* < 0.001, *****P*<0.0001, ns, not significant.

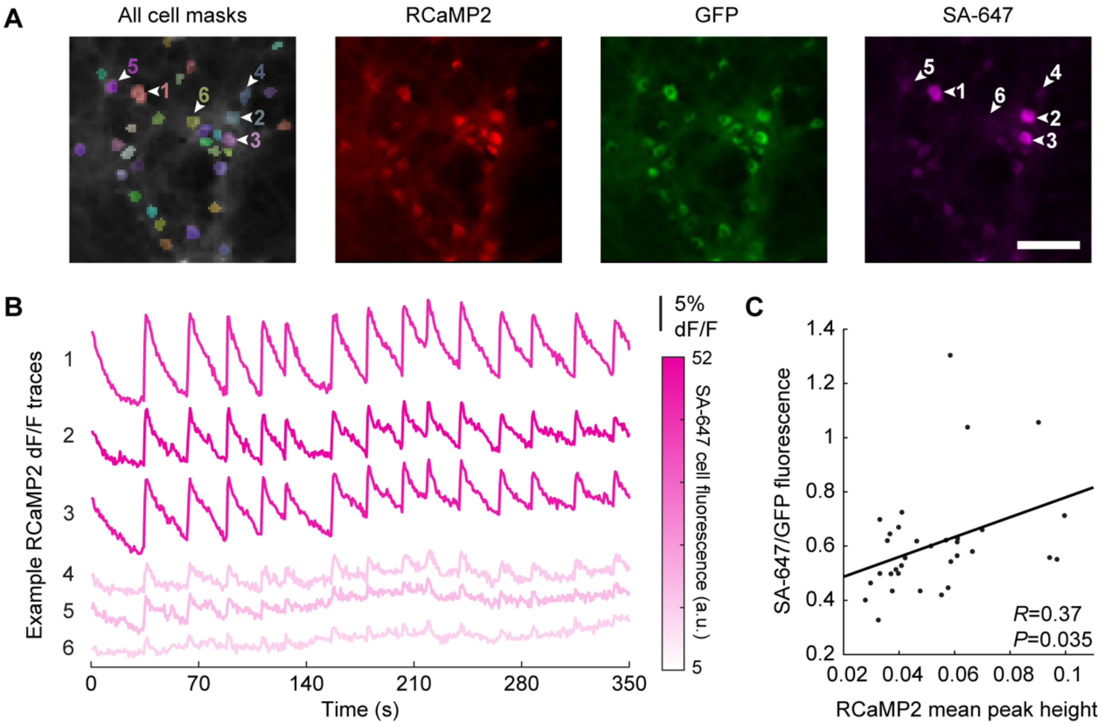


**Extended Data Figure 6. Simultaneous RCaMP2 imaging and CaST labeling in neurons.**
**A)** Example FOV images of neurons co-infected with AAV2/1-Synapsin-RCaMP2 and AAV2/1-Synapsin-CaST viruses, following mild stimulation (50% media change) and 50 µM biotin treatment for 30 minutes. Post-hoc RCaMP2 and CaST labeling is shown for all identified cell masks during RCaMP2 imaging (shown as colored overlays to the left). Numbered arrows indicate locations from which traces were extracted for panel **B**. Scale bar, 100 µm.
**B)** RCaMP2 dF/F fluorescence traces of example neurons from panel **A**, during a ~5-minute recording following treatment. Traces are colored according to their SA-647 cell fluorescence intensity value.
**C)** Scatter plot showing a linear correlation between SA-647/GFP fluorescence ratio calculated for each GFP+ neuron detected, and the mean peak height during the RCaMP2 recording for each cell ($n$ = 33 cells; two-tailed Pearson's correlation coefficient $R$ = 0.37, $P$ = 0.035).

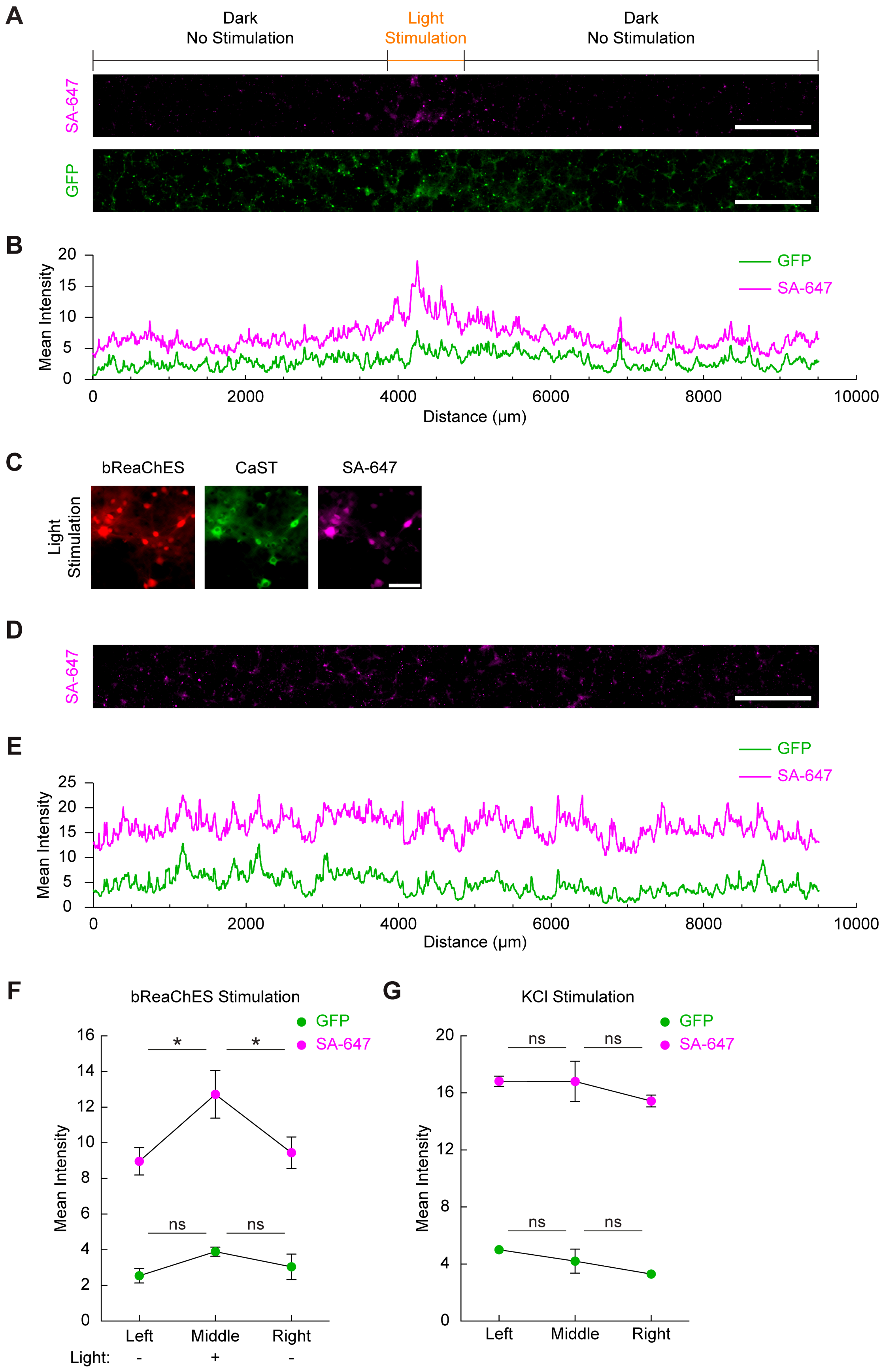
A
Dark
No Stimulation
Light
Stimulation
Dark
No Stimulation
SA-647
GFP
B
Mean Intensity
GFP
SA-647
Distance (µm)
C
bReaChES
CaST
SA-647
Light
Stimulation
D
SA-647
E
Mean Intensity
GFP
SA-647
Distance (µm)
F
bReaChES Stimulation
GFP
SA-647
Mean Intensity
Left
Middle
Right
Light:
-
+
-
G
KCl Stimulation
GFP
SA-647
Mean Intensity
Left
Middle
Right

**Extended Data Figure 7. CaST specificity using targeted optogenetic stimulation.**
**A)** Example fluorescence images for neurons co-infected with an excitatory opsin, AAV2/1-Synapsin-mCherry-P2A-bReaChES, and AAV2/1-Synapsin-CaST. 50 μM biotin and orange light was delivered for 30 minutes through a ~ 1mm wide slit to the bottom of the culture dish. 50 μM APV and 20 μM NBQX were added at the time of light stimulation to reduce synchronized neuron firing. Scale bar, 1000 μm.
**B)** Quantification of the mean SA-647 and GFP fluorescence intensity, averaged vertically across the entire FOV images shown in panel **A**.
**C)** Example zoom-in images of the light-stimulated region in panel **A** showing neurons co-expressing bReaChES and CaST. Scale bar, 100 μm.
**D)** Example SA-647 image for CaST-expressing neurons with whole dish KCl treatment as a non-spatial control. Scale bar, 1000 μm.
**E)** Quantification of the mean SA-647 and GFP fluorescence intensity, averaged vertically across the entire FOV shown in panel **D**.
**F)** Mean SA-647 and GFP fluorescence intensity, averaged vertically and binned across 1 mm horizontal sections to the left, middle, or right of the light stimulation gap (For SA-647, $n$ = 3 FOVs per condition; Left versus Middle: $P$ = 0.0281; Middle versus Right: $P$ = 0.0449, Šídák's post-hoc multiple comparison's test following a 2-way ANOVA, $F_{2,4}$ = 12.94, $P$ = 0.0179) (For GFP, $n$ = 3 FOVs per condition; Left versus Middle: $P$ = 0.1645; Middle versus Right: $P$ = 0.4352, Šídák's post-hoc multiple comparison's test following a 2-way ANOVA, $F_{2,4}$ = 3.536, $P$ = 0.1305).
**G)** Same analysis as in panel **F**, except for a whole dish KCl stimulation non-spatial control (For SA-647, $n$ = 3 FOVs per condition; Left versus Middle: $P$ = 1.0; Middle versus Right: $P$ = 0.6731, Šídák's post-hoc multiple comparison's test following a 2-way ANOVA, $F_{2,4}$ = 0.9006, $P$ = 0.4754) (For GFP, $n$ = 3 FOVs per condition; Left versus Middle: $P$ = 0.7328; Middle versus Right: $P$ = 0.6682, Šídák's post-hoc multiple comparison's test following a 2-way ANOVA, $F_{2,4}$ = 2.446, $P$ = 0.2023). Data are plotted as mean ± s.e.m. in **F** and **G**. *$P$<0.05, ns, not significant.

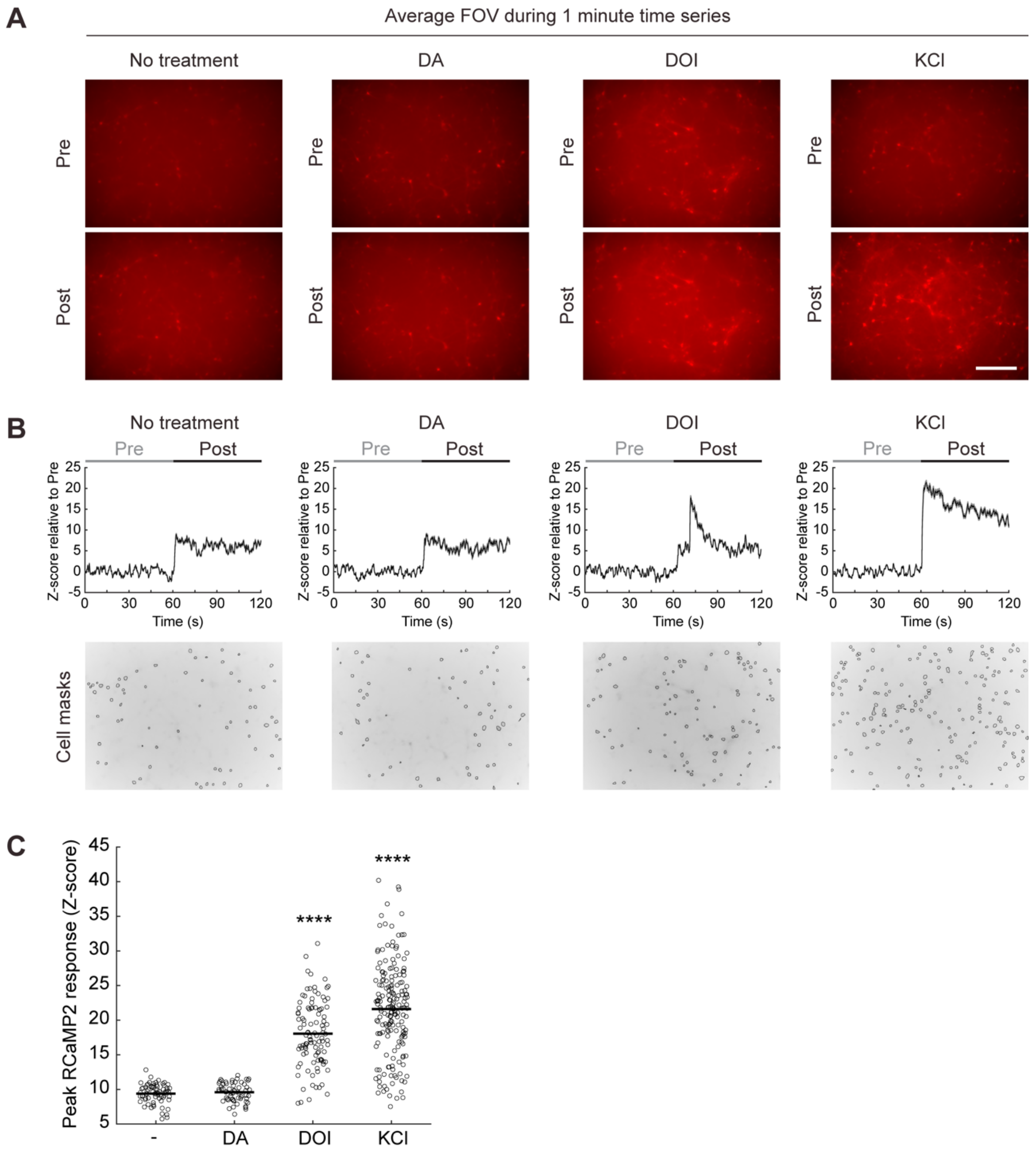


**Extended Data Figure 8. RCaMP2 calcium imaging during various drug applications.**
**A)** Average FOV images of neurons infected with AAV2/1-Synapsin-RCamp2, taken from the entire RCaMP2 baseline recording ("Pre") or post-treatment recording ("Post"). Each recording was 1 minute long. Neurons were treated with 50 μM biotin and vehicle, 10 μM dopamine (DA), 10 μM DOI, or 30 mM KCl for 30 minutes. Scale bar, 300 μm.
**B)** Top: average pre- and post-treatment fluorescence traces for all identified neurons in the FOV (Z-scored relative to the baseline "pre" period for each cell). Bottom: cell masks identified for each FOV shown in panel **A**. Data are plotted as mean ± s.e.m. with the shaded regions indicating the s.e.m.
**C)** Peak neuron responses during the post-treatment recordings. Only DOI and KCl treatment drove a larger peak RCaMP2 response compared to the vehicle control (*n* = 69 cells for vehicle,

62 cells for DA, 104 cells for DOI, and 189 cells for KCl treatment; No KCl versus DOI: *P* = 2.3e-13; No KCl versus KCl: *P* = 2.3e-13, Tukey's post-hoc multiple comparison's test following a 1-way ANOVA, $F_{3,420}$ = 152.4, *P* = 7.8e-67). ****$P$<0.0001.

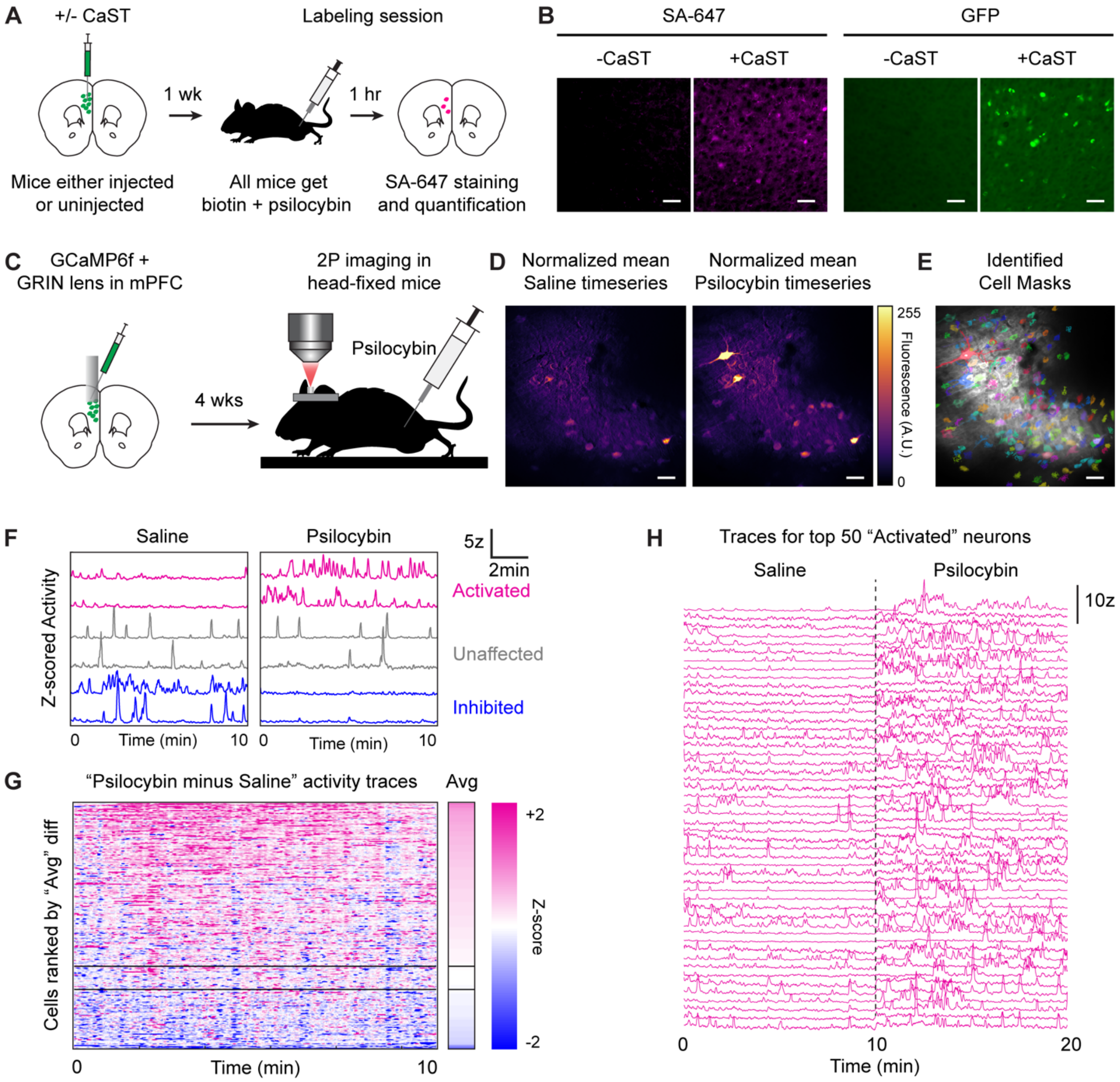


**Extended Data Figure 9. Controls and validation for in vivo CaST labeling.**

**A)** Schematic of control and experimental conditions. Both wildtype mice not expressing CaST, and wildtype mice injected with CaST in mPFC, were treated with 24 mg/kg biotin + 3 mg/kg psilocybin for 1 hour, and then sacrificed for histology.

**B)** Example FOVs of wildtype mice not expressing CaST (-CaST) or expressing CaST (+CaST) that were injected with biotin and psilocybin. The -CaST control was replicated across two uninjected mice.

**C)** Schematic for using real-time imaging to identify psilocybin-activated neurons in the mPFC. AAV5-CaMKIIa-GCaMP6f was injected into mPFC, and a 1mm diameter GRIN lens was implanted. 4 weeks later, mice were imaged head-fixed under a 2P microscope during an IP injection of 5 ml/kg saline or 3 mg/kg psilocybin.

**D)** Background-subtracted mean images of the FOV during a 10-minute saline recording session (left), and a 10-minute psilocybin recording session (right). Active neurons are displayed as warmer pixel colors. Experiment was replicated in two mice.

**E)** Mean 2P FOV image of the combined saline and psilocybin recordings with all identified neuron masks shown as colored overlays.
**F)** Example Z-scored fluorescence traces of neurons activated by psilocybin (magenta), unaffected by psilocybin (gray), or inhibited by psilocybin (blue), compared to the saline recording. Each recording was 10 minutes long.
**G)** "Psilocybin minus Saline" activity traces were calculated by subtracting the baseline saline Z-scored trace from the psilocybin Z-scored trace for each neuron. The resulting 10-minute-long trace representing the difference is plotted for each neuron in the heatmap to the left, and the average difference for each neuron is plotted as the heatmap to the right labeled "Avg" ($N$ = 254 cells from 2 mice). Neurons are plotted ranked by the highest to lowest average Z-score difference. The two horizontal bars represent the thresholds for defining "Activated" versus "Inhibited" neurons (>+0.05 = "Activated", <-0.05 = "Inhibited").
**H)** The activity traces for the top 50 ranked "Activated" neurons from panel **G** are shown during the saline and psilocybin recordings (separated by a dashed vertical line). All scale bars, 50 μm.

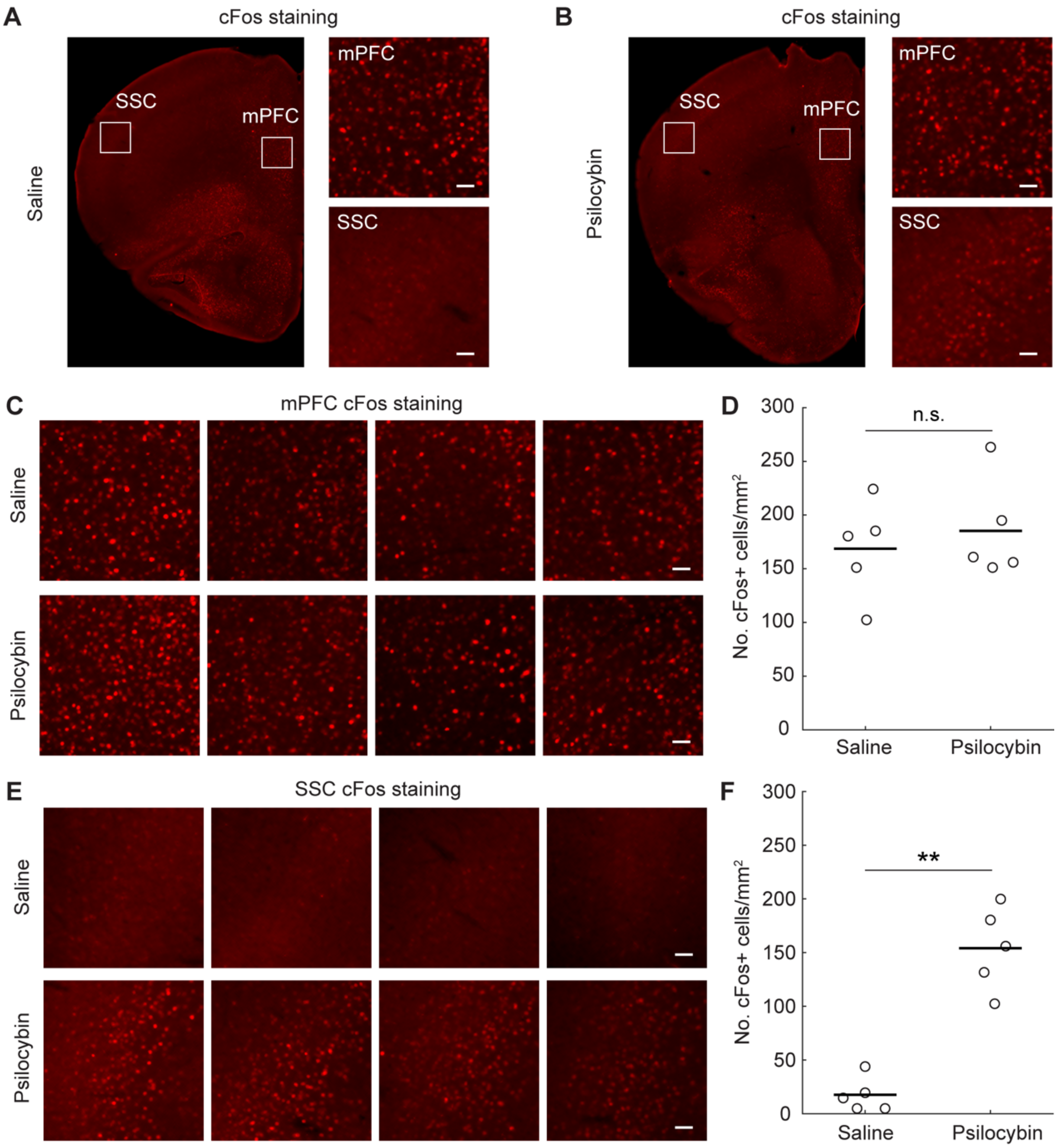


**Extended Data Figure 10. cFos-only staining in psilocybin- versus saline-injected mice.**
**A,B)** 2x (left) and 10x (right) images of mouse brain slices stained for cFos 1 hour after injection with saline vehicle (**A**) or 2 mg/kg psilocybin I.P. (**B**). White boxes on 2x images show location where the 10x images were taken in either the mPFC or SSC.
**C)** An additional 4 example FOVs taken from different mice showing cFos staining in the mPFC after saline or psilocybin injection as described in panel **A**.
**D)** Mean number of cFos+ neurons/mm$^2$ in the mPFC counted from mice injected with either saline or psilocybin ($n$ = 5 mice each condition; $P$ = 0.73, U = 10.50, two-tailed Mann-Whitney U test).
**E)** An additional 4 example FOVs taken from the same mice as in panel **C**, except showing cFos staining in the SSC after saline or psilocybin injection.
**F)** Mean number of cFos+ neurons/mm$^2$ in the SSC counted from mice injected with either saline or psilocybin ($n$ = 5 mice each condition; $P$ = 0.0079, U = 0, two-tailed Mann-Whitney U test). All scale bars, 50 μm. **$P$<0.01, ns, not significant.
