## Supplemental Figs 1-3, Table 1, Note 1 for "Rapid, biochemical tagging of cellular activity history in vivo"

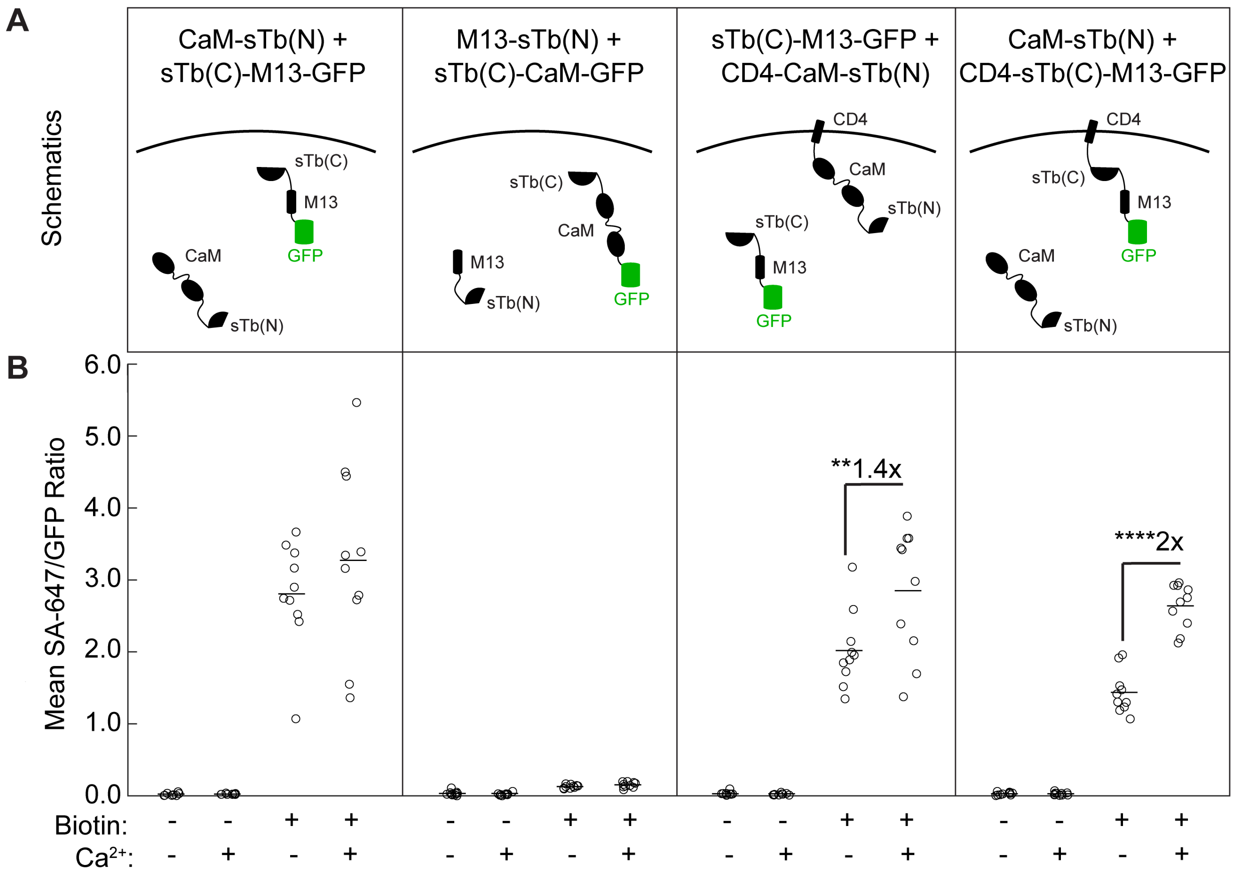


**Supplementary Figure 1. Comparison of different CaST versions.**

**A)** Schematics of 4 different initial CaST versions tested, with unique conformations and subcellular targeting of different components.

**B)** HEK cells were transfected with the constructs shown for each version in panel **A**. Cells were treated with ± 50 μM biotin and ± $Ca^{2+}$ (5 mM $CaCl_2$ and 1 μM ionomycin) for 30 minutes. Cells were then fixed, washed, stained with SA-647, and imaged. The FOV averages of the SA-647/GFP fluorescence ratio per cell were calculated across all conditions. Version 1 did not exhibit a significant interaction between biotin and $Ca^{2+}$ treatment ($n$ = 10 FOVs per condition; Šídák's post-hoc multiple comparison's test following a 2-way ANOVA, $F_{1,36} = 0.99$, $P = 0.33$). Version 2 resulted in low SA-647 labeling across all conditions. Version 3 exhibited a significant interaction between biotin and $Ca^{2+}$ treatment with a significant ~1.4-fold ± $Ca^{2+}$ SBR ($n$ = 10 FOVs per condition; $P = 0.0056$, Šídák's post-hoc multiple comparison's test following a 2-way ANOVA, $F_{1,36} = 6.64$, $P = 0.014$). Version 4, with CaM-sTb(N) + CD4-sTb(C)-M13-GFP, also exhibited a significant interaction between biotin and $Ca^{2+}$ treatment with a significant ~2-fold ± $Ca^{2+}$ SBR under these conditions ($n$ = 10 FOVs per condition; $P$ = 7.1e-14, Šídák's post-hoc multiple comparison's test following a 2-way ANOVA, $F_{1,36} = 78.19$, $P$ = 1.5e-10).
$^{**}P < 0.01$, $^{****}P < 0.0001$.

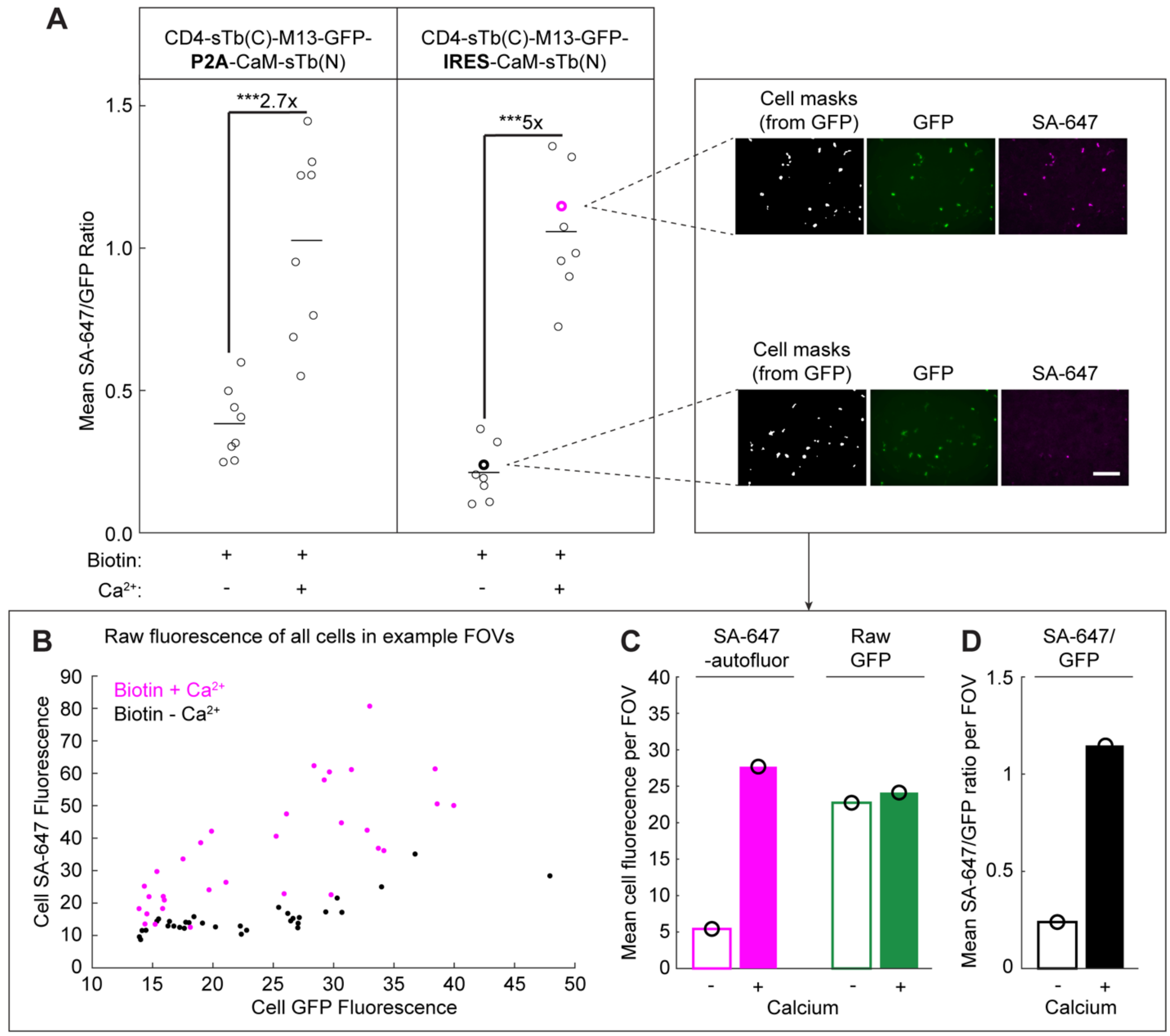


**Supplementary Figure 2. P2A vs IRES comparison and analysis pipeline in HEK cells.**
**A)** HEK cells were transfected with either a P2A and IRES version of the final CaST design, and cells were treated with biotin ± $Ca^{2+}$ for 30 minutes as in panel **A**. The P2A version exhibited a ± $Ca^{2+}$ SBR of 2.7x (*n* = 8 per condition; *P* = 3.1e-4, U = 37, two-tailed Mann-Whitney U test). The IRES version exhibited a ± $Ca^{2+}$ SBR of 5x (*n* = 8 per condition; *P* = 1.6e-4, U = 36, two-tailed Mann-Whitney U test). Example FOV images from CaST-IRES group with all identified cell masks are shown on the right. Scale bar, 300 μm.
**B)** Scatter plot of the mean SA-647 versus mean GFP fluorescence calculated for every GFP+ cell detected across the two FOVs highlighted in panel **A.**
**C)** Magenta: the mean SA-647 cell fluorescence (minus the background autofluorescence) for all GFP+ cells shown in the two highlighted FOVs in panel **B**. The background autofluorescence in the SA-647 image was calculated as the mean value of the entire image excluding pixels corresponding to cell masks. Green: the mean GFP cell fluorescence calculated for all GFP+ cells shown in the two highlighted FOVs in panel **B**.
**D)** The mean SA-647/GFP ratio of all GFP+ cells shown in panel **B**, calculated by dividing each cell's background-subtracted SA-647 fluorescence value by its GFP fluorescence value.
***$P$ < 0.001.

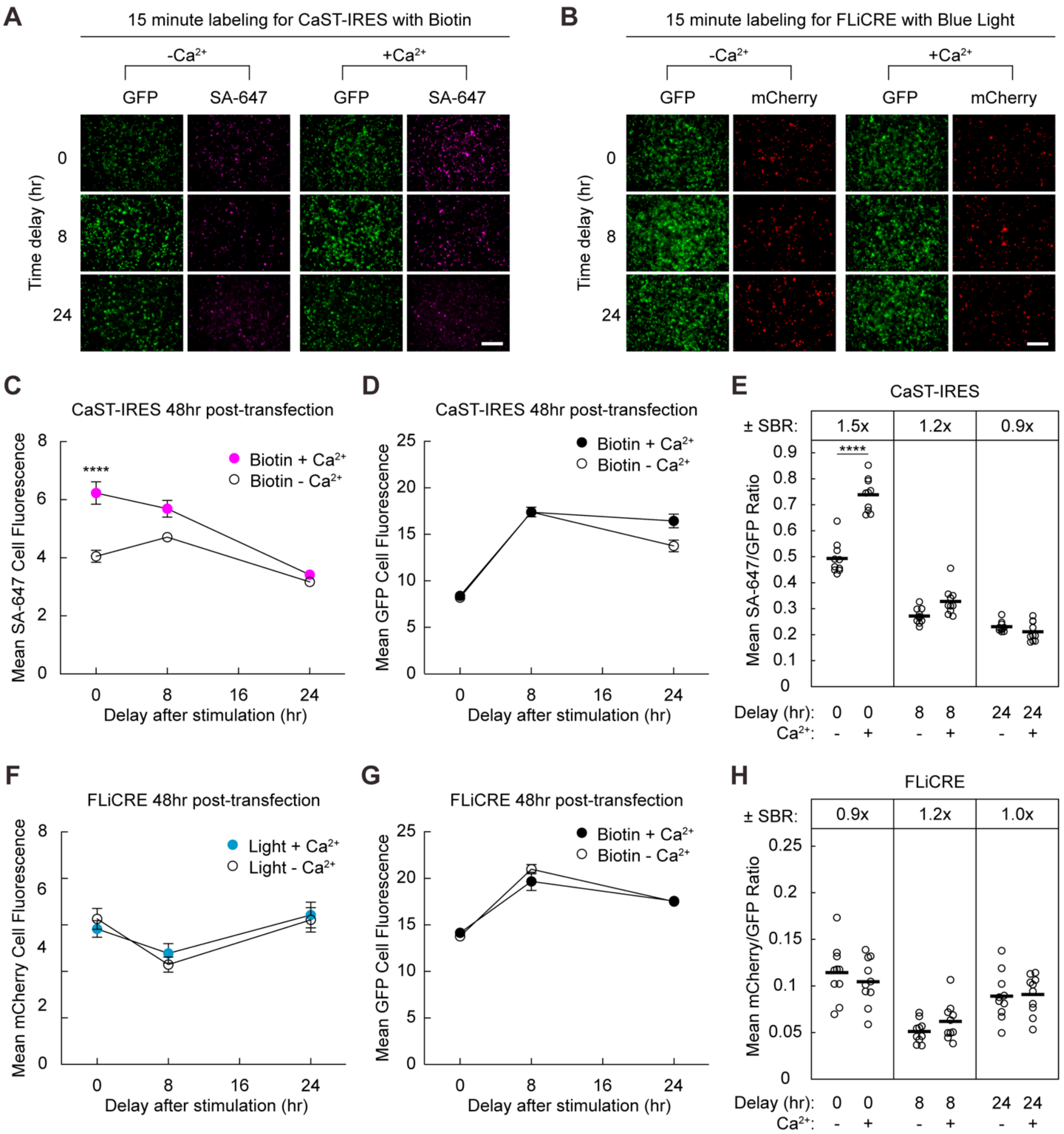


**Supplementary Figure 3. Comparison of CaST to FLiCRE at higher expression levels.**
**A,B)** Example FOVs for CaST and FLiCRE 48 hours post-transfection experiments with 0, 8, and 24 hours delay after 15 minutes stimulation**.** Scale bar, 300 µm.
**C)** For CaST-IRES, cells were transfected with CaST-IRES and were incubated 48 hours prior to stimulation. The FOV average of the SA-647 cell fluorescence was calculated following a variable delay period after biotin ± $Ca^{2+}$ stimulation shown in panel **A** (*n* = 10 FOVs per condition; *P* = 3.0e-7, Šídák's post-hoc multiple comparison's test following a 2-way ANOVA, $F_{2,54}$ = 8.670, *P* = 0.0005).
**D)** The FOV average of the GFP cell fluorescence for CaST-IRES shown in panel **A** (*n* = 10 FOVs per condition).

**E)** The ± $Ca^{2+}$ SBR of normalized reporter expression for 0, 8, and 24 hours after stimulation for CaST-IRES. For CaST, the SA-647 fluorescence is divided by the GFP fluorescence (*n* = 10 FOVs per condition; *P* = 1.1e-5, U = 0, two-tailed Mann-Whitney U test).
**F)** For FLiCRE, cells were transfected with FLiCRE components and were incubated 48 hours prior to stimulation. The FOV average of the UAS::mCherry cell fluorescence was calculated following a variable delay period after light ± $Ca^{2+}$ stimulation shown in panel **B** (*n* = 10 FOVs per condition, Šídák's post-hoc multiple comparison's test following a 2-way ANOVA, $F_{2,54}$ = 0.5587, *P* = 0.5752).
**G)** The FOV average of the UAS::mCherry cell fluorescence for FLiCRE shown in panel **B** (*n* = 10 FOVs per condition).
**H)** The ± $Ca^{2+}$ SBR of normalized reporter expression for 0, 8, and 24 hours after stimulation for FLiCRE. For FLiCRE, the UAS::mCherry fluorescence is divided by the GFP fluorescence (*n* = 10 FOVs per condition). Data are plotted as mean ± s.e.m. in **C**-**D**, **F**-**G**.
****$P$ < 0.0001.

**Supplementary Table 1. Plasmids used or generated in this study**.

| Insert Name | Description | Vector-Promoter | Access |
|---|---|---|---|
| CaM-V5-sTb(N) | sTb(N) half of CaST for HEK cell expression | AAV-CMV | Addgene #219779 |
| CD4-sTb(C)-M13-GFP | sTb(C) half CaST for HEK cell expression | AAV-CMV | Addgene #219780 |
| sTb(C)-CaM-GFP | Alternative unused variant of sTb(C) fusion | AAV-CMV | Available upon request |
| M13-V5-sTb(N) | Alternative unused variant of sTb(N) fusion | AAV-CMV | Available upon request |
| CD4-CaM-V5-sTb(N) | Alternative unused variant of sTb(N) fusion | AAV-CMV | Available upon request |
| sTb(C)-M13-GFP | Alternative unused variant of sTb(C) fusion | AAV-CMV | Available upon request |
| CD4-sTb(C)-M13-GFP-P2A-CaM-V5-sTN | CaST-P2A for HEK cell expression | AAV-CMV | Addgene #219781 |
| CD4-sTb(C)-M13-GFP-IRES-CaM-V5-sTb(N) | CaST-IRES for HEK cell expression | AAV-CMV | Addgene #219782 |
| CD4-MKII-hLOV1-TEVcs(ENLYFQ/M)-Gal4 | FLiCRE TF for HEK cell expression | AAV-CMV | Addgene #163026 |
| mCherry | FLiCRE reporter for HEK cell expression | AAV-UAS | Addgene #135457 |
| GFP-CaM-uTEVp | FLiCRE protease for HEK cell expression | AAV-CMV | Addgene #163028 |
| CaM-V5-sTb(N) | sTb(N) half of CaST for neuron expression | AAV-Synapsin | Addgene #219783 |
| CD4-sTb(C)-M13-GFP | sTb(C) half of CaST for neuron expression | AAV-Synapsin | Addgene #219784 |
| RCaMP2 | Red Calcium indicator for neuron expression | AAV-Synapsin | Available upon request |
| mCherry-p2A-bReaChES-HA | Red shifted excitatory opsin for neuron expression | AAV-Synapsin | Available upon request |

**Supplementary Note 1. Considerations for in vivo biotin delivery during CaST labeling.**

Biotin is actively transported through the blood brain barrier through native transporters via its carboxyl group. A previous study used PET radioactive ligand binding experiments to record biotin trafficking in vivo in mice using a minimally modified $^{11}$C-biotin (biotin labeled with radionuclide carbon-11). Importantly, the carboxyl group of $^{11}$C-biotin is untouched, so that it retains the same structural and biochemical properties as native biotin. This study performed PET imaging following $^{11}$C-biotin injection I.V. and showed a sustained presence of $^{11}$C-biotin in the brain over the duration of 1 hour. Once transported into the brain, it remains trapped there[1], until it is eventually excreted through the blood and then urine over the course of several hours ($t_{90\%}$ = 12 hours for a 10mg/kg I.P. dose of $^{14}$C-biotin[2]).

Based on these studies, we reason that the optimal time window for CaST labeling should be during the first initial hour after biotin injection. Thus, we recommend injecting mice with 24 mg/kg biotin I.P. (the dose used for in vivo TurboID[3] and BioID[4] labeling) immediately prior to the experimental stimulus. In the case where the stimulus is a drug also being delivered I.P., the biotin and drug can be co-administered at the same time. If the stimulus is a behavioral event expected to drive a prolonged elevated change in neuronal activity (e.g., exposure to a stressor, or a change in metabolic environment), the biotin can be administered ~5 minutes prior to the event to allow for I.P. transport time to the brain.

In addition, given that biotin remains trapped in the brain, we recommend immediately sacrificing and perfusing the mice after the experimental labeling paradigm, to reduce accumulating background biotinylation. We note that a negative control experiment should always be performed, where mice are injected with biotin but not exposed to the drug or behavioral stimulus thought to induce neuronal activity. CaST labeling quantification and selection of positively-labeled neurons should always be performed in comparison to the background labeling determined in this negative control condition.

Future studies are needed to determine whether repeated administration of biotin (e.g., repeated I.P. injections or food/water-supplemented biotin) would enable CaST labeling over longer periods of time.


1. Bongarzone, S. et al. Imaging biotin trafficking in vivo with positron emission tomography. *Journal of medicinal chemistry* **63**, 8265-8275 (2020).
2. Lee, H.-M., Wright, L.D. & McCormick, D.B. Metabolism of carbonyl-labeled 14C-biotin in the rat. *The Journal of Nutrition* **102**, 1453-1463 (1972).
3. Kim, K.-e. et al. Dynamic tracking and identification of tissue-specific secretory proteins in the circulation of live mice. *Nature Communications* **12**, 5204 (2021).
4. Uezu, A. et al. Identification of an elaborate complex mediating postsynaptic inhibition. *Science* **353**, 1123-1129 (2016).
